# The Cdc14 phosphatase controls resolution of recombination intermediates and crossover formation during meiosis

**DOI:** 10.1101/571083

**Authors:** Paula Alonso-Ramos, David Álvarez-Melo, Katerina Strouhalova, Carolina Pascual-Silva, George B. Garside, Meret Arter, Teresa Bermejo, Rokas Grigaitis, Rahel Wettstein, Marta Fernández-Díaz, Joao Matos, Marco Geymonat, Pedro A. San-Segundo, Jesús A. Carballo

## Abstract

Meiotic defects derived from incorrect DNA repair during gametogenesis can lead to mutations, aneuploidies and infertility. Coordinated resolution of meiotic recombination intermediates is required for crossover formation, ultimately necessary for accurate completion of both rounds of chromosome segregation. Numerous master kinases orchestrate the correct assembly and activity of the repair machinery. Although much less is known, reversal of phosphorylation events in meiosis must also be key to coordinate the timing and functionality of repair enzymes. Cdc14 is an evolutionarily conserved phosphatase required for the dephosphorylation of multiple CDK1 targets. Mutations that inactivate this phosphatase lead to meiotic failure, but until now it was unknown if Cdc14 plays a direct role in meiotic recombination. Here, we show that elimination of Cdc14 leads to severe defects in the processing and resolution of recombination intermediates, causing a drastic depletion of crossovers when other repair pathways are compromised. We also show that Cdc14 is required for correct activity and localization of the Holliday Junction resolvase Yen1/GEN1. We reveal that Cdc14 regulates Yen1 activity from meiosis I onwards, and this function is essential for crossover resolution in the absence of other repair pathways. We also demonstrate that Cdc14 and Yen1 are required to safeguard sister chromatid segregation during the second meiotic division, a late action that is independent of the earlier role in crossover formation. Thus, this work uncovers previously undescribed functions of Cdc14 in the regulation of meiotic recombination.

## Introduction

Meiotic recombination is initiated by the conserved Spo11 transesterase, which introduces numerous DNA Double-Strand Breaks (DSBs) into the genome (Keeney et al. 1997). Association of single-strand DNA binding proteins, including RPA, Rad51 and the meiosis specific recombinase, Dmc1, with the resected DSB ends promotes strand invasion into the intact homologous non-sister chromatid template, which culminates in the formation of a displacement loop (D-loop). Nascent D-loops can be processed through the Synthesis-Dependent Strand Annealing (SDSA) repair pathway to generate non-crossover (NCO) repair products (Bishop 2006). Alternatively, stabilization of the D-loop followed by second-end capture gives rise to even more stable structures, such as double Holliday Junction (dHJ) intermediates. In budding yeast meiosis, dHJs are most frequently resolved into crossovers (COs) through the combined action of Mlh1-Mlh3 (MutLγ) and Exo1 (Zakharyevich et al. 2010; Zakharyevich et al. 2012; Sanchez et al. 2020). A second class of COs arises from the resolution of recombination intermediates via the Structure-Selective Endonucleases (SSEs) Mus81/Mms4, Slx1/Slx4 and Yen1 (de los Santos et al. 2001; Jessop and Lichten 2008; Oh et al. 2008; Matos et al. 2011; Arter et al. 2018). Finally, Sgs1, the ortholog of the Bloom’s syndrome helicase (BLM), can also promote dissolution of JMs via the Sgs1-Top3-Rmi1 (STR) complex, which would exclusively generate NCOs products. In addition, multiple rounds of strand exchange over a stabilized D-loop, can give rise to three and four interconnected duplexes, also known as multichromatid joint molecules (mc-JMs) (Oh et al. 2007). Further, both DSB-ends might also participate in multi-invasion events on two or three chromatids, creating heavily branched DNA structures (Piazza et al. 2017). These complex DNA species must be efficiently processed to prevent the risk of becoming potential hazards for the genome integrity due to their capacity to form aberrant crossovers, or other forms of toxic repair products. Correct processing of all types of JMs requires the orchestrated action of a set of helicases, topoisomerases and endonucleases, capable of dislodging multi-branched DNA assemblies. The BLM/Sgs1, in addition to be involved in SDSA repair, plays a prominent role in eliminating aberrant “off-pathway” JMs, including mc-JMs (Jessop and Lichten 2008; Oh et al. 2008; De Muyt et al. 2012; Copsey et al. 2013; Xaver et al. 2013; Grigaitis et al. 2018).

In budding yeast, it is not fully clear which enzymatic complexes take over the function of removing any unprocessed branched DNA intermediates outside prophase I, due to the existence of numerous SSEs capable of processing such substrates (Machin 2020; San-Segundo and Clemente-Blanco 2020). Upon initiation of *NDT80-*dependent transcription, and activation of the budding yeast polo-like kinase Cdc5, cells abandon the pachytene stage concomitantly with a surge of enzymatic activity that culminates in the resolution of ZMM-dependent and ZMM-independent dHJs to give rise to COs and some NCO products (Clyne et al. 2003; Sourirajan and Lichten 2008). Cdc5 phosphorylation is required to stimulate the function of a set of SSEs, most notoriously Mus81-Mms4 (Matos et al. 2011; Wild et al. 2019). The Mus81-Mms4 complex acts on branched DNA substrates that have not been resolved by the canonical ZMM-dependent Class-I CO pathway (De Muyt et al. 2012; Zakharyevich et al. 2012). On the other hand, Yen1^GEN1^, a conserved member of the Rad2/XPG family of SSEs, appears to act later in meiosis, despite the fact that Yen1 is considered a prototypical HJ resolvase (Ip et al. 2008). In meiotic cells, this discrepancy is easily explained since the activity of Yen1 is negatively regulated by CDK1-dependent phosphorylation, preventing the activation of the nuclease during prophase I (Matos et al. 2011). Elimination of CDK-phosphorylation sites in Yen1, turns on its enzymatic activity prematurely and promotes its nuclear localization. Therefore, counteracting the phosphorylation of Yen1 generates the fully active form of the nuclease (Matos et al. 2011; Arter et al. 2018). In mitotic cells, dephosphorylation of Yen1 is carried out by the Cdc14 phosphatase (Blanco et al. 2014; Eissler et al. 2014). Activation of Yen1 during anaphase by Cdc14 allows the resolution of persistent repair intermediates that otherwise would impose a physical impediment to chromosome segregation. It is currently unknown whether Cdc14 also activates Yen1 in meiosis (Matos et al. 2011; Zakharyevich et al. 2012; Grigaitis et al. 2020). Furthermore, it is possible that Cdc14 dephosphorylates other substrates important for the timely resolution of different types of JMs (Grigaitis et al. 2020).

Cdc14 is a well-conserved dual-specificity phosphatase that has been defined as a key component in the reversal of CDK phosphorylation during exit from mitosis (Visintin et al. 1998). In budding yeast, Cdc14 activity is essential and cells lacking this phosphatase remain arrested at the end of mitosis (Culotti and Hartwell 1971; Hartwell et al. 1973). Additionally, Cdc14 regulates transcriptional silencing at the rDNA and other repeated sequences (Machin et al. 2006; Clemente-Blanco et al. 2009; Clemente-Blanco et al. 2011). Several pathways regulate Cdc14 localization and activity. The RENT complex retains Cdc14 at the nucleolus until anaphase through its interaction with the Cif1/Net1 protein (Shou et al. 1999; Visintin et al. 1999). Cdc14 is released from the nucleolus by the sequential action of two pathways. The FEAR (Cdc <u>F</u>ourteen <u>E</u>arly <u>A</u>naphase <u>R</u>elease) pathway promotes the early release of Cdc14 through the phosphorylation of Cif1/Net1 and Cdc14 (Yoshida and Toh-e 2002; Visintin et al. 2003; Azzam et al. 2004; Queralt et al. 2006; Rahal and Amon 2008; Rodriguez-Rodriguez et al. 2016; Zhou et al. 2021). Later in anaphase, the Mitotic Exit Network (MEN) keeps Cdc14 in its released state, allowing the dephosphorylation of additional substrates, important for the full termination of mitosis and cytokinesis (Stegmeier et al. 2002; Visintin et al. 2008).

In addition to curtailing CDK activity at the end of mitosis, Cdc14 plays critical roles promoting chromosome segregation (Hartwell and Smith 1985). In *S. cerevisiae*, it is required for the correct segregation of the ribosomal gene array (rDNA) and telomeres (D’Amours et al. 2004; Sullivan et al. 2004; Torres-Rosell et al. 2004). Lack of transcriptional silencing at these regions prevents the loading of condensins at the rDNA and subtelomeric regions (D’Amours et al. 2004; Wang et al. 2004; Clemente-Blanco et al. 2011), which leads to unresolved linkages, intertwined sister chromatids, and non-disjunction during mid-anaphase. Notably, a number of functions have been allocated to Cdc14 before its anaphase-triggered release. In particular, Cdc14 has been involved in completion of replication of late-replicating regions, such as the rDNA locus and other parts of the genome (Dulev et al. 2009). Additionally, anaphase-independent release of Cdc14 has been observed upon induction of DNA damage; generation of DSBs promotes the transitory release of Cdc14 from the nucleolus, targeting the SPB component Spc110 for dephosphorylation (Villoria et al. 2017).

Cdc14 is highly conserved and orthologs have been identified in other organisms, including fission yeast, nematodes, insects, and vertebrates (Mocciaro and Schiebel 2010). The human genome contains three Cdc14 paralogs, hCDC14A, hCDC14B and hCDC14C (Li et al. 1997; Rosso et al. 2008). Depletion of Cdc14A, which is required for mitotic CDK inactivation, leads to defects in centrosome duplication, mitosis and cytokinesis (Trautmann and McCollum 2002). In vertebrates, CDC14B exits the nucleolus in response to DNA damage (Bassermann et al. 2008; Mocciaro et al. 2010), a process conserved in fission yeast (Diaz-Cuervo and Bueno 2008), and more recently also observed in budding yeast (Villoria et al. 2017). Thus, in addition to playing important roles in the DNA Damage Response (DDR) (Bassermann et al. 2008) it appears that both Cdc14A and Cdc14B may be required for efficient repair of damaged DNA (Mocciaro et al. 2010).

Cdc14 is also critically required for the completion of meiosis (Sharon and Simchen 1990). In budding yeast, Cdc14 is released from the nucleolus in two waves; the first during anaphase I and the second in anaphase II (Buonomo et al. 2003; Marston et al. 2003; Yellman and Roeder 2015). Contrary to mitotic cells, Cdc14 release from the nucleolus requires the essential function of the FEAR pathway, but MEN appears to be dispensable (Kamieniecki et al. 2005; Pablo-Hernando et al. 2007; Attner and Amon 2012). Additionally, some components of the MEN pathway, such as Cdc15, have functionally differentiated to fulfil a role in spore morphogenesis (Kamieniecki et al. 2005; Pablo-Hernando et al. 2007). Curiously, inactivation of the FEAR pathway in meiosis allows exclusively one round of nuclear division to take place, culminating with the formation of asci carrying two diploid spores (dyads) (Kamieniecki et al. 2000; Zeng and Saunders 2000; Buonomo et al. 2003; Marston et al. 2003; Kamieniecki et al. 2005). Premature activation of FEAR blocks spindle assembly during meiosis, a process normally averted by PP2A^Cdc55^ (Queralt et al. 2006; Bizzari and Marston 2011; Kerr et al. 2011; Nolt et al. 2011). On the other hand, inactivation of Cdc14 function by means of FEAR mutants, or by employing conditional temperature-sensitive alleles of *CDC14*, impairs the second round of chromosome segregation (Buonomo et al. 2003; Marston et al. 2003). Furthermore, the absence of the phosphatase creates chromosome entanglements, a problem that can be reverted by deleting *SPO11* (Marston et al. 2003). Occasionally, in the absence of Cdc14, the second meiotic division takes place over non-tetrapolar meiosis II spindles and, subsequently, these cells carry out aberrant chromosome segregation (Bizzari and Marston 2011). These interesting observations prompted the identification of another critical function of Cdc14 at the meiosis I to meiosis II transition, which is the licensing of SPB re-duplication/half-bridge separation by asymmetrical enrichment of Cdc14 on a single SPB during anaphase I (Fox et al. 2017).

Here, we describe a previously undefined role of Cdc14 in budding yeast meiosis. We reveal that Cdc14 is involved in ensuring correct meiotic recombination by gradually implementing Yen1 activity following Cdc5 activation. We show that, at anaphase I, Yen1 already shows a prominent nuclear localization promoted by Cdc14. Yen1 reaches its maximum activity during meiosis II, and Cdc14 is strictly required for such activation. A constitutively active form of Yen1, termed Yen1^ON^, which is insensitive to CDK downregulation, compensates for Cdc14 deficiency, suggesting that Cdc14 normally reverses CDK phosphorylation in Yen1 during meiosis. Unexpectedly, Cdc14 also promotes Yen1 activity in *CDC20*-depleted cells, allowing JM processing and CO formation when other repair pathways are absent. Thus, Yen1 plays a more prominent role during meiotic recombination than originally anticipated, which can be manifested even before meiosis I division is normally completed. Notably, Yen1 complements other endonucleases whose activities are mainly constrained to the time-window between the end of prophase I and before anaphase I. We propose that Yen1, controlled by Cdc14, implements its resolvase function when other enzymatic activities begin to decline outside of their optimum operative time-window by the end of prophase I, and that Yen1 covers a wider gap extending its action until the end of the second meiotic division.

## Results

### Identification of a separation-of-function allele of *CDC14* defective in sporulation

Reversal of CDK1 phosphorylation is carried out in budding yeast by the conserved Cdc14 phosphatase. This essential protein can only be studied utilizing conditional *cdc14* alleles that fully or partially impair its functionality. Thermosensitive mutants have been widely used in the past for mitotic and meiotic studies. However, meiotic recombination is sensitive to variations of temperature (Borner et al. 2004), thus we considered using other types of conditional alleles to further examine the role of Cdc14 in meiosis. During the course of a different study, we generated the *cdc14-HA* allele carrying an endogenous C-terminal tagged version of *CDC14* with three copies of the hemagglutinin epitope (HA). The *cdc14-HA* strain was highly defective for sporulation, displaying only 10% of asci containing one or two spores (Fig 1A), and lacking dityrosine autofluorescence, an indicator of spore wall maturation (Briza et al. 1986) (Fig 1B & F and figure S2B). Unlike other temperature-sensitive alleles of *cdc14* used in many studies (see introduction), mitotic growth was not affected in diploid cells carrying the *cdc14-HA* allele incubated at different temperatures (Fig 1C). Thus, we conclude that *cdc14-HA* cells cannot efficiently complete sporulation despite their apparently normal mitotic growth and may be a useful tool to explore additional aspects of Cdc14 meiotic function.

**Figure 1.**
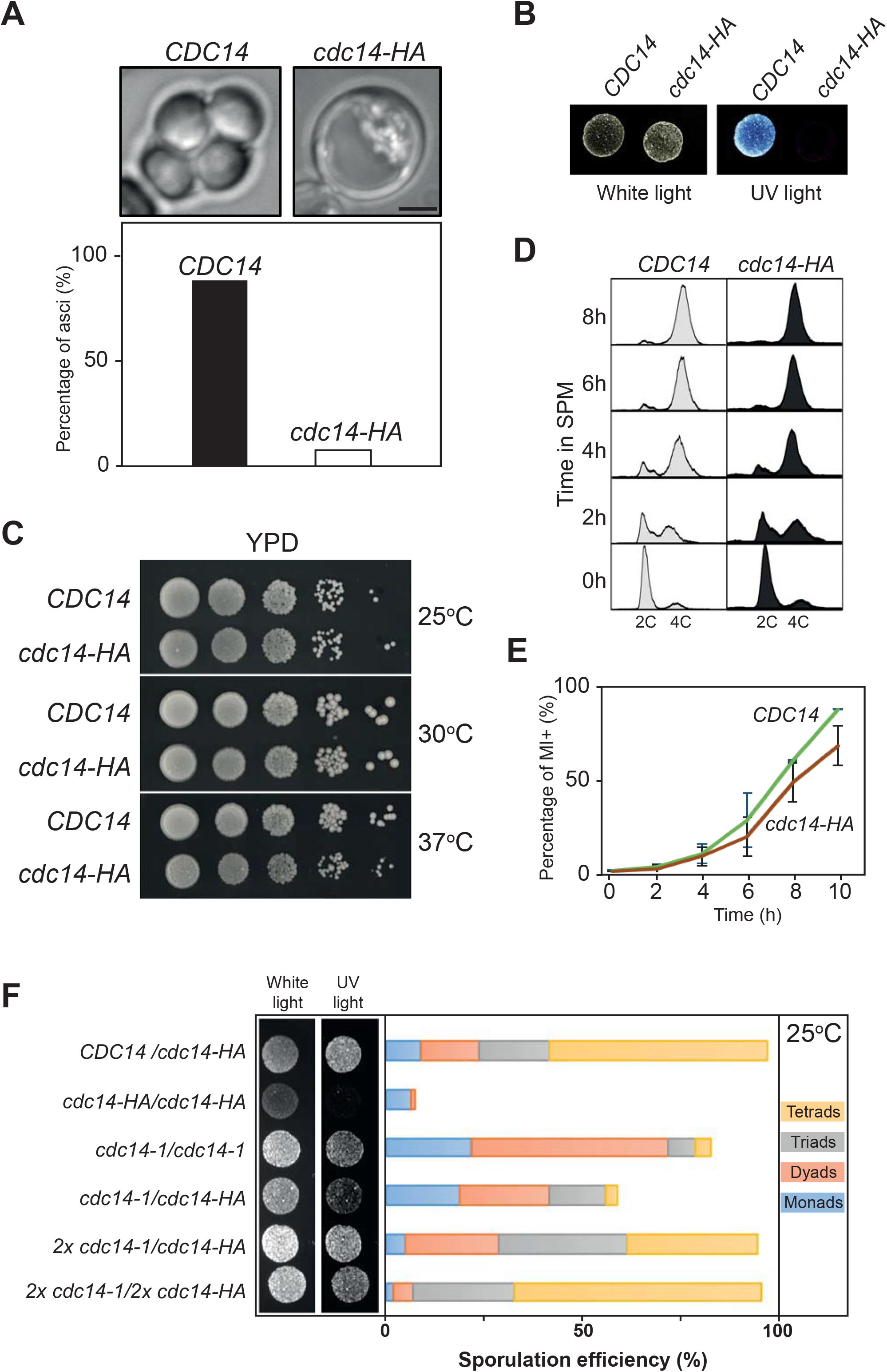
*cdc14-HA* behaves as a specific sporulation-deficient separation-of-function allele of *CDC14*. (A) Homozygous *cdc14-HA* diploids do not form asci containing spores under standard conditions for sporulation. (B) *cdc14-HA* diploid cells divide mitotically but lack di-tyrosine autofluorescence induced by UV light after several days of incubation in SPM media at 30°C. (C) The *cdc14-HA* allele does not perturb normal growth conditions when cultivated at 25°C, 30°C and 37°C in rich media (YPD). (D) FACS analysis of DNA content shows that *cdc14-HA* cells complete meiotic DNA replication with identical kinetics as *CDC14* diploid cells. (E) *cdc14-HA* cells undergo meiotic nuclear divisions at subtly slower kinetics, and reduced frequencies, than *CDC14* cells. Error bars represent the SEM over the mean values plotted. (F) Di-tyrosine autofluorescence of different mutant combinations as well as the control strains grown and sporulated on plates at 25°C. *cdc14-1* homozygous diploids (JCY2353) sporulate at high efficiency under semi-permissive temperature forming preferentially tetrads whereas *cdc14-HA* homozygous diploids (JCY840) do not sporulate. Combinations, and variable copy number, of the mutant genes can rescue the sporulation defect to different degrees (JCY2365/JCY2354/JCY2356).

### Distinct sporulation-defective alleles of *CDC14* block meiotic progression at different stages

To determine the cause(s) of the sporulation defect of the *cdc14-HA* mutant we analyzed the progression of the meiotic program by studying the kinetics of meiotic DNA replication and nuclear divisions. During the mitotic cell cycle, released Cdc14 dephosphorylates replication factors, such as Sld2 and Dpb2 (Bloom and Cross 2007), and the lack of Cdc14 activity leads to problems during termination of DNA replication (Dulev et al. 2009). However, FACS analysis of DNA content revealed that S-phase progression occurred with nearly identical kinetics in both control and *cdc14-HA* strains (Fig 1D), suggesting that meiotic S-phase is not altered in *cdc14-HA*. Next, meiotic divisions were monitored in synchronous cultures of *CDC14* and *cdc14-HA*. We found that *cdc14-HA* cells displayed slightly slower and less efficient kinetics of nuclear divisions; nevertheless, most *cdc14-HA* cells exited the one nuclei stage, although with a ∼60 min delay compared to wild-type cells (Fig 1E; Fig. S1A). Previous work has shown that most meiotic cells in *cdc14* temperature-sensitive mutants are blocked with two nuclei, due to the inability to assemble fully functional tetrapolar spindles at meiosis II (Schild and Byers 1980; Sharon and Simchen 1990; Marston et al. 2003; Bizzari and Marston 2011). However, we noticed that the *cdc14-HA* mutant progressed beyond the two-nuclei stage in a high proportion of cells (Fig S1B-C), distinguishing this *cdc14-HA* allele from temperature-sensitive mutants used in other studies.

Next, we generated an additional meiosis-specific allele of *CDC14* by replacing its endogenous promoter with the *CLB2* promoter, which is strongly repressed in meiotic cells (Lee and Amon 2003). In this case, a 3HA tag was located at the N-terminal region of the protein. Cells expressing *P_CLB2_-3HA-CDC14* (hereafter, *cdc14-md*, for <u>m</u>eiotic <u>d</u>epletion) recapitulated the sporulation defect of *cdc14-HA* (Fig 3G). The most striking difference was that although the first meiotic nuclear division took place in *cdc14-md* as it occurs in *cdc14-HA*, most cells remained binucleated in *cdc14-md* even at late time points in meiosis (Fig S1A). Thus, the terminal meiotic phenotype of *cdc14-md* appears to be different from that of *cdc14-HA*.

### Cdc14 protein levels are depleted during meiosis in *cdc14-HA* cells

We next used western blot to analyze Cdc14 protein levels during meiosis in *CDC14, cdc14-md* and *cdc14-HA* strains. The *cdc14-HA* mutant displayed reduced levels of the tagged protein, even at time zero, prior to meiotic induction, and they were further reduced after 6-8 hours into meiosis (Fig 2A and Fig S2A-B). Depletion of Cdc14 was more effective in the *cdc14-md* allele, while in *cdc14-HA* it occurred more gradually (Fig 2A). This is not related to delayed entry into the meiotic program, since all strains underwent meiotic S-phase with similar kinetics as determined by FACS analysis (Fig 2A). To directly ascertain if Cdc14 protein levels varied from mitotically dividing cells to meiotic cells, we also analyzed extracts from exponentially growing cultures compared to extracts prepared from cultures immediately after meiotic induction (Fig 2B). The wild-type strain showed similar amounts of Cdc14 protein in both mitotic and meiotic cultures. The levels of Cdc14-HA were lower in mitotic cells compared to the wild type, and they were further reduce upon meiotic entry. On the other hand, Cdc14 protein levels in the mitotically dividing *cdc14-md* mutant were similar to the wild type, but they showed a drastic depletion with meiosis onset (Fig 2B). Increasing the copy number of different *cdc14* alleles, as well as overproduction of the Cdc14-HA protein restored sporulation to a large extent (Fig 1F and Fig S2C-D). These results indicate that the reduction of Cdc14 phosphatase levels, but not its full depletion, is causing the meiotic phenotype in the hypomorphic *cdc14-HA* mutant. On the other hand, the almost complete absence of the protein throughout the entire meiotic process likely provokes the more stringent block in nuclear divisions observed in *cdc14-md* cells. Thus, both alleles are useful tools to investigate different meiotic events that could be regulated by Cdc14.

**Figure 2.**
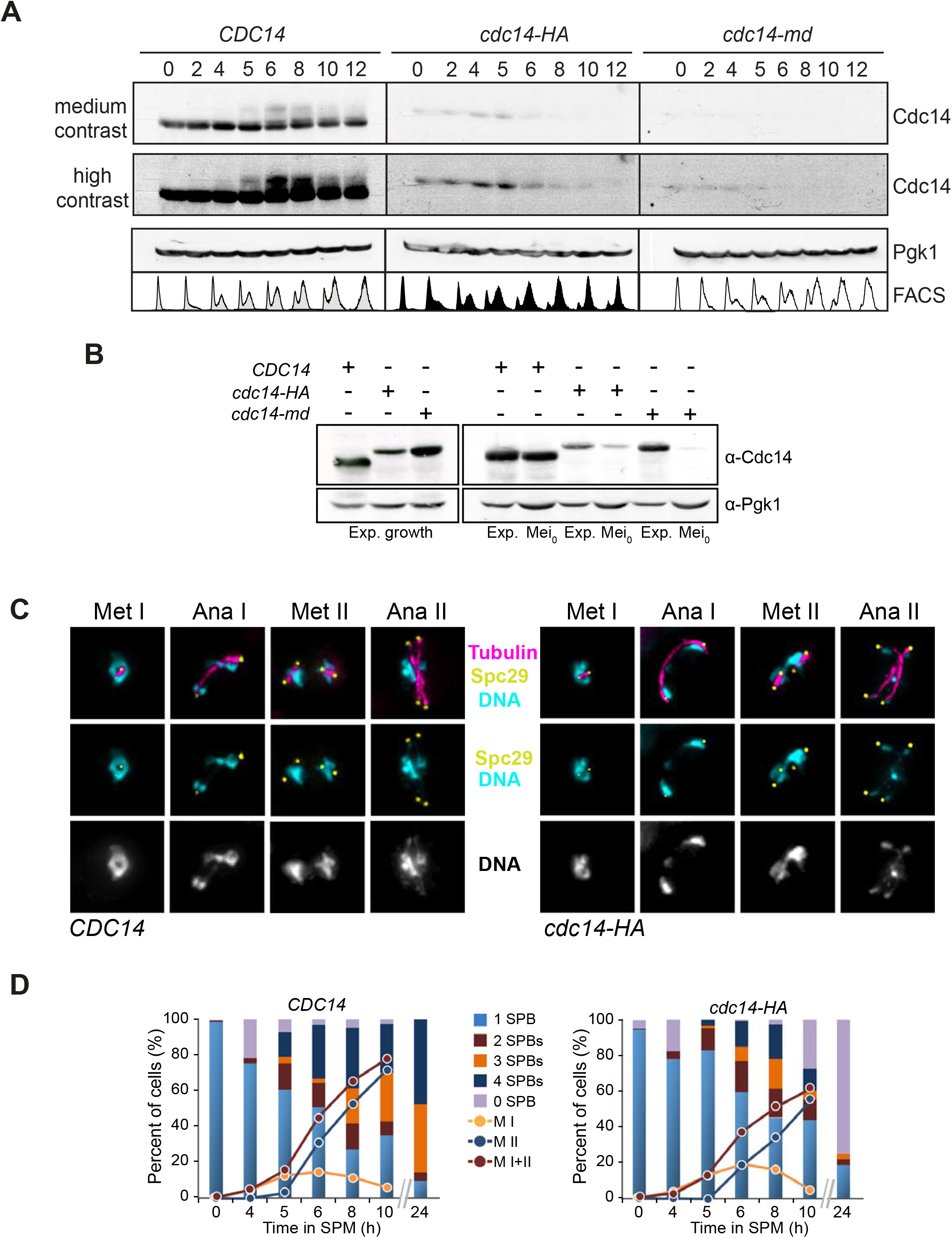
Diminished Cdc14 protein levels in *cdc14-HA* meiotic cells compromise SPB integrity following meiotic divisions. (A) Dynamics of Cdc14 protein levels throughout meiosis in synchronous time-courses for *CDC14* (JCY2232), *cdc14-HA* (JCY2231), and *cdc14-md* (JCY2389). Higher contrast was applied to visualize residual protein levels for all three strains. Immunodetection of Pgk1 was used as loading control. FACS histograms for each time-point are depicted at the bottom to show the degree of synchrony reached in each culture. (B) Variability of protein levels in mitotic versus meiotic cells in different *CDC14* alleles. Exponentially growing cells show similar Cdc14 levels except for *cdc14-HA*, which has lower overall protein levels (left panel). Direct comparison between the three alleles *CDC14* (JCY2232); *cdc14-HA* (JCY2231) and *cdc14-md* (JCY2389) in exponentially growing mitotic cells (Exp.) and meiotic cultures immediately after they were transferred to SPM (Mei_0_). Loading control for each lane was determined using Pgk1. (C) Visualization of Spc29-CFP at SPBs, tubulin, and DNA, in wild-type (JCY892) and *cdc14-HA* (JCY893) cells fixed at different stages of meiosis at 30°C. (D) *cdc14-HA* cells develop meiosis I and II spindles with near wild-type kinetics. Lack of spore formation in *cdc14-HA* meiotic cells parallels the loss of SPB’s structural integrity.

### SPB integrity is compromised in *cdc14-HA* cells after the second meiotic division

As mentioned above, a great proportion of *cdc14-HA* cells eventually passed the two-nuclei stage observed as terminal meiotic phenotype in *cdc14-md* (this study) and *cdc14* temperature-sensitive mutants (Schild and Byers 1980; Sharon and Simchen 1990; Marston et al. 2003). Thus, we monitored the kinetics and morphology of the spindle pole bodies (SPBs) and spindles during the first and the second meiotic divisions (Fig 2C-D). We observed that by eight hours into meiosis, *CDC14* and *cdc14-HA* cultures contained around 75% and 60% of cells with more than one SPB, respectively. Among the population of cells that showed multiple SPBs, 56% of wild-type cells presented more than two; therefore, they had initiated meiosis II. However, only 40% of *cdc14-HA* cells displayed more than two SPBs, confirming a slight reduction of cells forming meiosis II spindles carrying four separated SPBs. This observation correlates with the slightly lower frequency of *cdc14-HA* cells containing more than two DAPI masses (Fig 2D; Fig S1B). As expected, after 24 hours of meiotic induction, the wild type showed over 90% of cells with three or four SPBs. In marked contrast, most *cdc14-HA* meiotic mutant cells lacked a single SPB, suggesting that, in the absence of Cdc14, the structural integrity of SPBs becomes compromised once both meiotic divisions take place (Fig 2D; Fig S3A). On the other hand, most cells of the *cdc14-md* mutant, contained only 2 SPBs for the entire length of the time course (Fig S3D), confirming earlier studies revealing a role for the phosphatase in SPB reduplication/half-bridge separation (Fox et al. 2017). Additionally, the average time taken to complete both meiotic divisions was not significantly different in the *cdc14-HA* mutant compared to the wild type (Fig S4A-C). Thus, we can conclude that, albeit with some difficulties, the *cdc14-HA* mutant is still capable of undergoing both meiotic divisions, but it is unable to form spores. On the other hand, the *cdc14-md* mutation, which produces a more drastic meiosis-specific depletion of Cdc14, prevents the separation of duplicated SPBs after the first division; therefore, these cells are unable to assemble proper tetrapolar spindles at meiosis II. Such behaviour resembles more closely the phenotype(s) previously described for other non-meiosis specific *cdc14* mutants (Marston et al. 2003; Fox et al. 2017).

### Impeded homolog separation and sister missegregation are common hallmarks of *cdc14* mutations in meiosis

We took advantage of the hypomorphic *cdc14-HA* allele, which allows us to study the second division in the absence, or with very little amount, of the Cdc14 protein to analyze chromosome segregation. In the *cdc14-HA* mutant, it was frequent to observe cells with more than four DAPI-stained masses late in meiosis (Fig S1B and Fig S3C). This might be due to impediments during chromosome segregation in the first and/or the second meiotic division. To test this possibility, we analyzed nuclear morphology, along with the meiotic spindle, in individual cells undergoing the first and second divisions (Fig 3Ai-v). Cells presenting meiosis I (MI) spindles with a length of ≥5μm were considered to be in late anaphase I, and therefore, two DAPI masses on each pole should be distinguishable (Fig 3Ai). On the other hand, cells presenting DAPI-stained bridges might reflect some form of chromosome entanglement (Fig 3Aii-iii; (Marston et al. 2003)). Wild-type cells showed a clear separation of nuclear content in 74% of anaphase I cells analyzed (Fig 3A). By contrast, in both *cdc14-HA* and *cdc14-md* mutants, only ∼20% of anaphase I cells displayed two fully resolved nuclei (Fig 3A). A large proportion (∼80%) of anaphase I mutant cells presented a chromosomal bridge, suggesting that despite having undergone MI spindle elongation correctly, some form of DNA entwining was not properly resolved.

**Figure 3.**
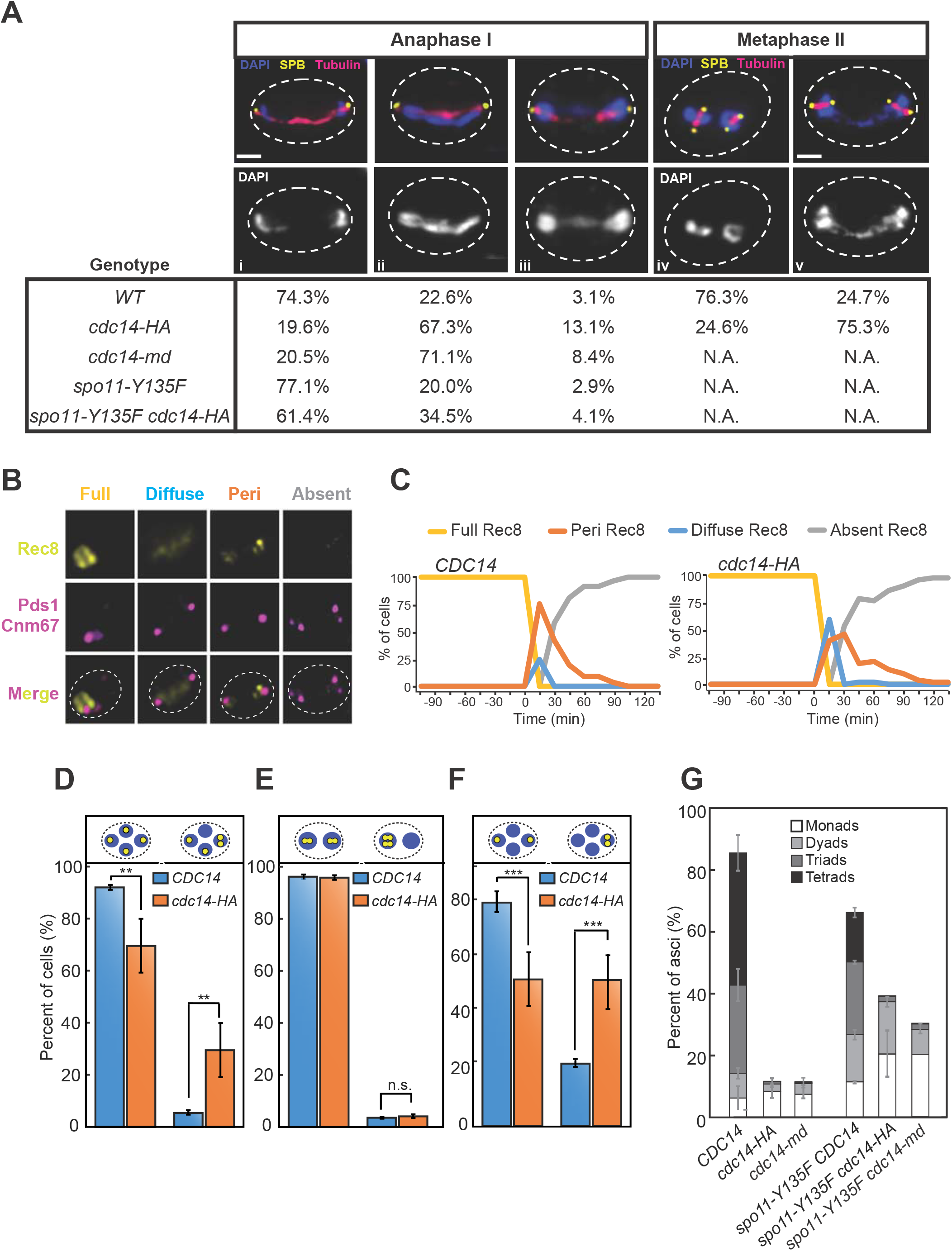
Missegregation of sister chromatids at MII in meiosis-deficient *cdc14* mutant cells. (A; i-iii) Frequency of cells presenting connected DAPI masses at anaphase I in wild type (JCY902), *cdc14-HA* (JCY904), *cdc14-md* (PRY132), *spo11* (PRY53) and *spo11 cdc14-HA* (PRY55). (A; iv-v) Frequency of cells presenting connected nuclei at metaphase II. Meiotic-deficient *cdc14-HA* mutants (GGY54) present higher frequencies of unresolved nuclear divisions at late anaphase I and at metaphase II than wild type (GGY53). (B) Representative images of distinct morphological patterns for Rec8-GFP in meiosis. Full: bright nuclear localization. Diffuse: faint staining within the stretched nucleus. Peri: Rec8-GFP only visible at peri-centromeric locations. Absent: no Rec8-GFP signal. Pds1-tdTomato and CNM67-tdTomato were used to follow Pds1 accumulation/degradation and SPB number/location. (C) Temporal distribution of the distinct morphological categories of Rec8-GFP shown in (B) for *CDC14* (JCY2406; n=57 cells) and *cdc14-HA* (JCY2404; n=52 cells). (D) Unequal distribution of homozygous chromosome II-linked GFP markers at *lys2* locus in *cdc14-HA* mutant cells (JCY2330) denotes increased chromosome missegregation in comparison with wild-type cells (JCY2331). (E) Similar distribution of homozygous *CENIV*-linked GFP markers at *trp1* locus in meiosis-deficient *cdc14-HA* mutant cells (JCY2286) in comparison with wild-type cells (JCY2284) denotes correct homolog disjunction in anaphase I. (F) Unequal distribution of heterozygous *CENIV-GFP* marker in meiosis-deficient *cdc14-HA* tetranucleated mutant cells (JCY2327) denotes increased sister chromatid missegregation in comparison with wild-type cells (JCY2326). Statistical significance of differences was calculated by two-tailed *t*-test, assuming unequal variances (\**p* < 0.05; \*\**p* < 0.01; \*\*\**p* < 0.001; \*\*\*\**p* < 0.0001; n.s.: not significant). (G) Sporulation defects in *cdc14-HA* (JCY844) or *cdc14-md* (JCY2376) can be partly alleviated (JCY840) by eliminating recombination (JCY2270/JCY2280/PRY151).

A subtle delay in cohesin removal during anaphase I onset was observed in a small proportion of *cdc14-HA* cells (Fig 3B-C and Fig S4D), which may be responsible for at least a fraction of these bridged DAPI masses (Marston et al. 2003). To test if these entanglements were eventually eliminated or persisted until the second meiotic division, we examined metaphase II cells and scored the configuration of their nuclei (Fig 3Aiv-v). The wild type presented over 76% of metaphase II cells with single individualized nuclei. By contrast, 75% of metaphase II cells in the *cdc14-HA* mutant showed DNA threads connecting both nuclear masses. As mentioned above, the severity of the SPB re-duplication/half-bridge separation phenotype in the *cdc14-md* strain precluded us to identify metaphase II cells with 4-SPBs in *cdc14-md* cells. We conclude that diminished levels of Cdc14 in meiosis results in the abnormal formation of nuclear bridges after the first meiotic division and that these bridges persist at least until metaphase II.

To determine whether the DNA bridges present at metaphase II in *cdc14-HA* mutant cells were due to problems arising during chromosome segregation, we constructed strains carrying fluorescent markers on specific chromosomes and followed their segregation (Straight et al. 1996). We made use of *tetO* repeats integrated at the interstitial *lys2* locus of chromosome II, together with the presence of the *tetR* repressor gene fused with GFP, which associates to these repeats (Michaelis et al. 1997; Katis et al. 2010). First, we analyzed strains labelled at chromosome II on both homologs to determine chromosome segregation fidelity in cells that had completed both meiotic divisions. We found that chromosome II segregated correctly in 95% and 78% of wild-type and *cdc14-HA* tetranucleated cells, respectively. The remaining 22% of *cdc14-HA* mutant cells displayed at least one of the four nuclei lacking a GFP focus (Fig 3D; Fig S5B type1, 2 and 3 only). We further analyzed the first and the second division separately using strains carrying *lacO* repeats at the *trp1* locus, which is linked to the centromere of chromosome IV (*CENIV*), and expressing the *lacI* repressor fused to GFP (Shonn et al. 2002) (Fig S5A). In order to assess homolog disjunction, we first examined binucleated cells homozygous for the *CENIV-GFP*, and we did not find any defect in the segregation of homologs during the first meiotic division in *cdc14-HA* (Fig 3E). Very similar results were previously reported for homozygous *CENV-GFP* markers in *cdc14-1* mutants (Bizzari and Marston 2011). Next, we analyzed sister chromatid segregation in the second meiotic division using heterozygous strains for the *CENIV-GFP* marker. In this case, 50% of tetranucleated *cdc14-HA* cells displayed problems segregating sister centromeres (Fig 3F). Taken all results together, we conclude that accurate sister chromatid segregation in meiosis requires unperturbed Cdc14 activity.

### Preventing recombination alleviates defective events in meiosis-specific *cdc14* mutants

It has been shown that elimination of meiotic DSBs alleviates problems with chromosome segregation in FEAR and *cdc14-1* mutants (Marston et al. 2003). Therefore, we considered that the presence of anaphase I nuclear bridges and sister chromatid missegregation in *cdc14-HA* cells could be, at least in part, originated from problems caused by defective recombination. If this were the case, these problems should be alleviated by eliminating meiotic recombination. To test this hypothesis, we combined the *cdc14-HA* allele with the *spo11-Y135F* mutation, which prevents DSB formation (Keeney et al. 1997). The absence of DSBs improved the separation of DAPI masses in anaphase I in both *cdc14-HA* and *cdc14-md* mutants (Fig 3A), indicating that recombination leads to the formation of anaphase I bridges in both meiosis-defective *cdc14* mutant strains. Accordingly, inactivation of Spo11 catalytic activity also improved sporulation in *cdc14-HA* and *cdc14-md* (Fig 3G). Thus, these results suggest that during meiosis Cdc14 plays a role in promoting accurate repair of Spo11-dependent DSBs.

### Meiotic recombination is impaired in *cdc14* mutants

To gain insight into the function of Cdc14 during meiotic recombination, we performed DNA physical assays using the well-characterized *HIS4LEU2* recombination hotspot (Fig 4A; (Cao et al. 1990; Schwacha and Kleckner 1994; Hunter and Kleckner 2001; Borner et al. 2004)) in wild type, *cdc14-HA* and *cdc14-md* strains. First, we measured total levels of DSBs, CO and NCOs products (Fig 4C-D). Levels of DSBs and COs were very similar in the three strains analyzed, albeit a small, not significant, reduction in NCO levels was observed in *cdc14-HA* and *cdc14-md* mutants (Fig 4C-D). Although a strong effect in the formation of recombination products was not perceived in *cdc14* mutant cells, shorter, or longer, lifespan of JMs could alter the ratio of NCO to CO to compensate for lower, or higher, efficiency in the processing of these intermediates (Martini et al. 2006; Oh et al. 2007; Shinohara et al. 2008). To address this possibility, we analyzed JM formation in the *ndt80Δ* mutant background (Allers and Lichten 2001). Accumulation of JMs takes place in *ndt80Δ* cells due to the lack of *CDC5*-dependent JM-resolution promoting activity (Sourirajan and Lichten 2008). DSBs, NCOs and JMs formed efficiently in the *cdc14-HA* mutant (Fig. 4E-F). Thus, under unchallenged conditions, Cdc14 appears to play little or no effect in the initiation and processing of meiotic recombination during prophase I, at least at the *HIS4LEU2* hotspot.

**Figure 4.**
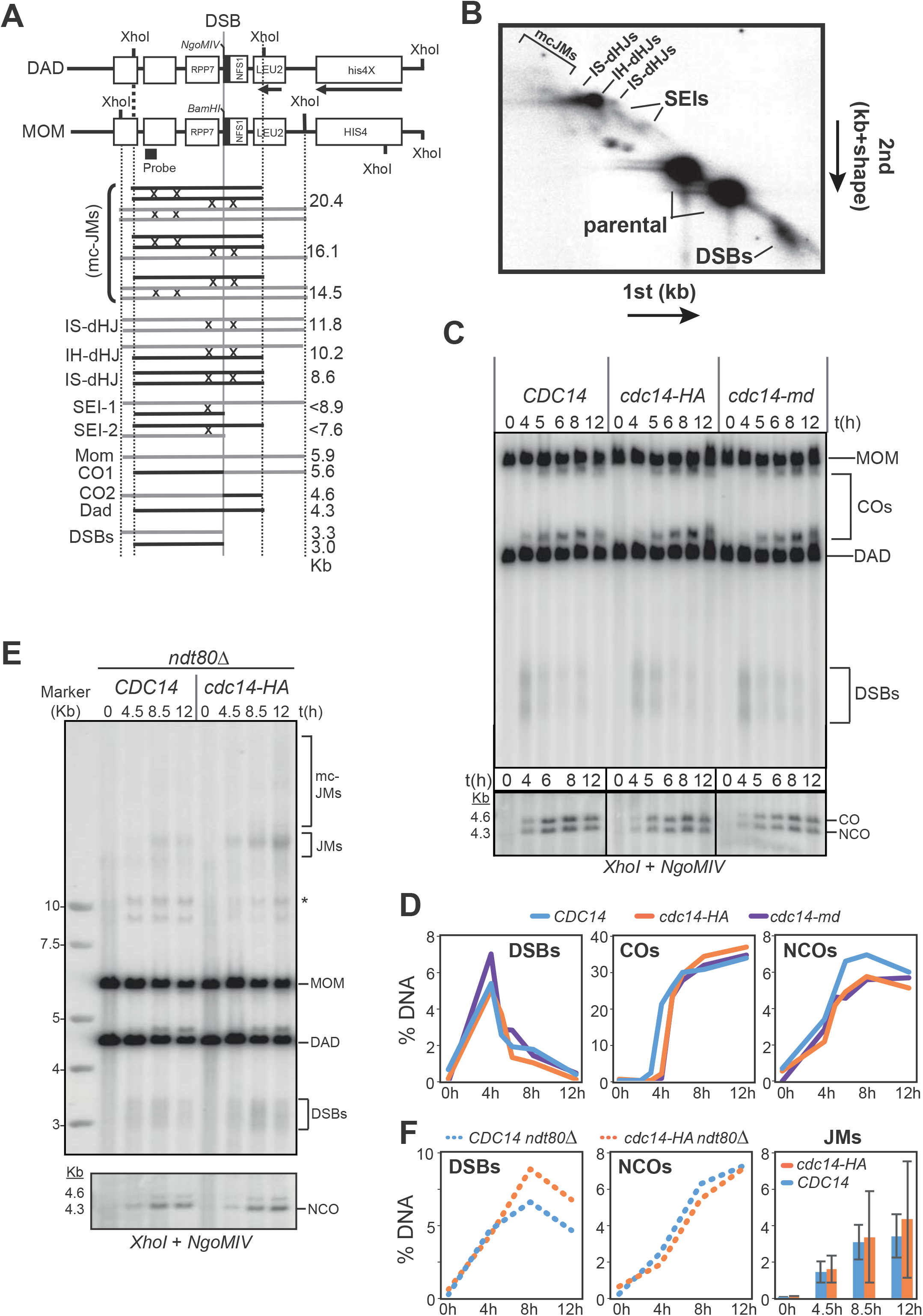
Analysis of meiotic recombination in meiotic-deficient *cdc14* mutants. (A) Schematic illustration of recombination analysis and intermediates at the *HIS4LEU2* hotspot (see Results for more details). (B) Schematic representation of DNA species commonly observed after analysing the *HIS4LEU2* hotspot by 2D-gel electrophoresis. (C; top) Representative 1D-gel Southern blot for analysis of DSBs and COs at the *HIS4LEU2* hotspot using *XhoI* as the restriction site. Wild-type strain (JCY2232), *cdc14-HA* (JCY2231), and *cdc14-md* (JCY2389). (C; bottom) Analysis of COs and NCOs at the *HIS4LEU2* hotspot using *XhoI* and *NgoMVI* in the same strains as in (C). (D) Quantification of DSBs, COs and NCOs from the Southern blots shown in (C). (E) Representative 1D-gel Southern blot image for analysis of DSBs, COs (top) and NCOs (bottom) at the *HIS4LEU2* hotspot in *CDC14 ndt8011* strain (JCY2390) and *cdc14-HA ndt8011* (JCY2385) strains. Asterisk indicates meiosis-specific non-characterized recombination products. (F) Quantification of DSBs, NCOs and JMs from the Southern blots shown in (E).

Next, we wanted to monitor if CO and/or NCO formation was affected in *cdc14* mutants when DSB repair is moderately compromised by eliminating one of the two main resolution pathways described in budding yeast, such as the Mus81/Mms4 pathway (de los Santos et al. 2001). Thus, we proceeded to evaluate the efficiency of JM resolution in a *mus81Δ ndt80Δ* mutant background. In order to obtain an accurate reading of repair product formation we first confirmed that total levels of JMs were similar in both *mus81Δ ndt80Δ* and *mus81Δ cdc14-HA ndt80Δ* mutant combinations (Fig 4B; Fig S6). To promote JM resolution in the absence of Ndt80, we took advantage of the widely-employed *CDC5*-inducible expression system (*CDC5-IN*), where the expression of the *CDC5*, which is required for the resolution of JMs into COs, can be induced at any point required in the meiotic time-course by the addition of β-estradiol (ES) to the media (Carlile and Amon 2008; Sourirajan and Lichten 2008). We added ES after 7 hours in meiosis and monitored CO and NCO formation. Induced expression of *CDC5* in the *mus81Δ* mutant efficiently led to the formation of COs only one hour after ES addition to the medium (Fig 5A-B); CO levels did not increase further after the 8-hour time-point. On the other hand, NCOs steadily increased reaching their maximum at 24h. In contrast, fewer COs were detected in the *mus81Δ cdc14-HA* mutant. NCOs levels were also lower in *mus81Δ cdc14-HA*, and total levels did not rise over the period when *CDC5* was expressed (Fig 5B). Thus, Cdc14 is required for efficient formation of COs and NCOs, at least when meiotic DSB repair takes place in the absence of the Mus81/Mms4-dependent resolution pathway.

**Figure 5.**
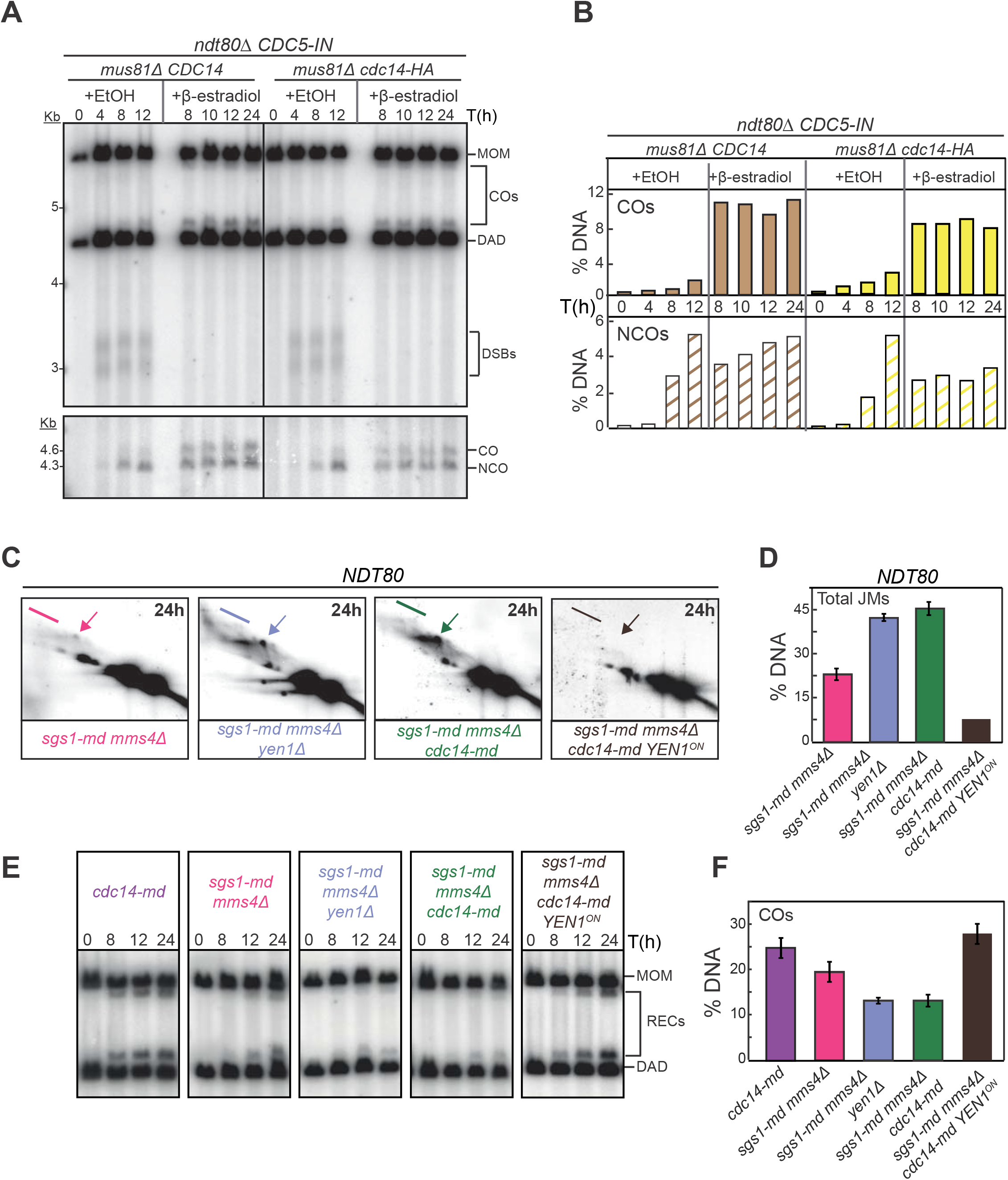
Analysis of recombination intermediates in *cdc14* mutants shows early and late accumulation of JMs. (A) Representative Southern blots depicting 1D-gels at the *HIS4LEU2* hotspot following induction of *CDC5* in prophase-arrested *ndt80Δ mus81Δ cdc14-HA* cells (JCY2440) leads to inefficient CO and NCO formation compared to *ndt80Δ mus81Δ* (JCY2442). (B) Quantification of COs and NCOs from Southern blots shown in (A). (C) Representative Southern blots depicting 2D-gels at the *HIS4LEU2* hotspot in *sgs1-md mms4Δ* (JCY2444), *sgs1-md mms4Δ cdc14-md* (JCY2446), *sgs1-md mms4Δ yen1Δ* (JCY2448) and *sgs1-md mms4Δ cdc14-md YEN1^ON^* (JCY2529) 24 h into meiosis. Arrows mark IH-dHJs and lines mark mc-JMs (see Fig 4A-B). (D) Quantification of total JMs in the strains shown in (C) from at least two independent gels. Error bars represent SEM over the calculated mean value. (E) Representative 1D-gel Southern blot images for analysis of crossovers at the *HIS4LEU2* hotspot for all strains shown in (C) and for *cdc14-md* (JCY2389). (F) Quantification of COs from at least three different image acquisitions like the one depicted in (E). Error bars represent SEM over the calculated mean value.

To confirm that Cdc14 plays a relevant role in the repair of recombination intermediates at least when certain resolution pathways are compromised, we decided to study the efficiency of repair in a *NDT80* background using 2D-gels that provide more precise information of the different DNA species involved during the JM resolution process (Fig 4B). To this end, we took advantage of the well-known negative effect on DSB-repair when depleting Sgs1 in combination with mutations in one or more of the SSEs, such as *mms4/mus81* and *yen1*/*slx1*/*slx4* (De Muyt et al. 2012; Zakharyevich et al. 2012). In this case, we used the *cdc14-md* allele, to prevent any residual activity of the Cdc14 phosphatase. As expected, elimination of *YEN1* in the *sgs1-md mms4Δ* background led to increased accumulation of unprocessed JMs compared to the *sgs1-md mms4Δ* double mutant (Fig 5C-D). Strikingly, after 24 hours in meiosis, the *sgs1-md mms4Δ cdc14-md* triple mutant also accumulated great levels of unresolved JMs compared to the *sgs1-md mms4Δ* double mutant (Fig 5C-D). Interestingly, the accumulation of unresolved JMs in *sgs1-md mms4Δ cdc14-md* was similar to that of the *sgs1-md mms4Δ yen1Δ* triple mutant (Fig 5C-D). Furthermore, CO levels at the *HIS4LEU2* hotspot were highly reduced in the *sgs1-md mms4Δ cdc14-md* strain; a phenotype that was nearly identical to that of *sgs1-md mms4Δ yen1Δ* (Fig 5E-F). Thus, in the absence of the Sgs1 and Mus81/Mms4-dependent pathways for processing recombination intermediates, the lack of the Cdc14 phosphatase leads to the same defects as the absence of the Yen1 resolvase, strongly suggesting that Cdc14 and Yen1 are involved in the same repair pathway during meiosis.

### Yen1 nuclear localization and activity during meiosis requires Cdc14 function

In mitotic cells, Yen1 localization is cytoplasmic when CDK activity is high, but release of Cdc14 during anaphase I reverts CDK-dependent phosphorylation on Yen1 promoting its nuclear entry and activity (Matos et al. 2011; Blanco et al. 2014; Eissler et al. 2014; Garcia-Luis et al. 2014). In meiotic cells, Yen1 subcellular localization is also regulated in a phosphorylation-dependent and cell-cycle stage-specific manner (Matos et al. 2011). To determine whether Cdc14 also regulates Yen1 subcellular distribution during meiosis, we examined nuclear localization of Yen1 in anaphase I cells, and we found that it was markedly reduced in *cdc14-HA* compared to the wild type (Fig 6A). We also tested if the enzymatic activity of Yen1 is controlled by Cdc14 during meiosis (Fig 6B-C). Samples were collected from synchronized meiotic cultures of wild-type and *cdc14-HA* strains carrying a functional 9MYC-tagged version of *YEN1* (Arter et al. 2018). From each time-point taken, Yen1 was immunoprecipitated from the meiotic extracts and the beads were incubated with a synthetic HJ substrate (Grigaitis et al. 2018). Active Yen1 should be able to cleave the HJ substrate whereas the phosphorylated inactive form of the nuclease would not (Blanco et al. 2014). In wild-type cells, Yen1 did not display significant nuclease activity during early stages of the meiotic time course prior to the accumulation of Cdc5 (Fig 6B), but approximately 2 hr after Cdc5 induction, we detected robust HJ processing, consistent with activation of Yen1 during meiotic divisions (Fig 6B-C; (Matos et al. 2011)). Notably, the resolvase activity displayed by Yen1 in the *cdc14-HA* mutant was extremely low for the duration of the whole time course (Fig 6B-C). Meiotic S-phase was completed with similar timing in wild-type and *cdc14-HA* cells suggesting that the initiation of the meiotic program occurred with similar kinetics in both strains (Fig 6D). Moreover, the Cdc5 protein (marking the exit from prophase I) was detected at the same time, and at the same levels, in both wild-type and *cdc14-HA* strains (Fig. 6B). Thus, these results strongly suggest that Cdc14 promotes nuclear accumulation of Yen1 during anaphase I and is required for its enzymatic activity at times when Cdc5 activity is high.

**Figure 6.**
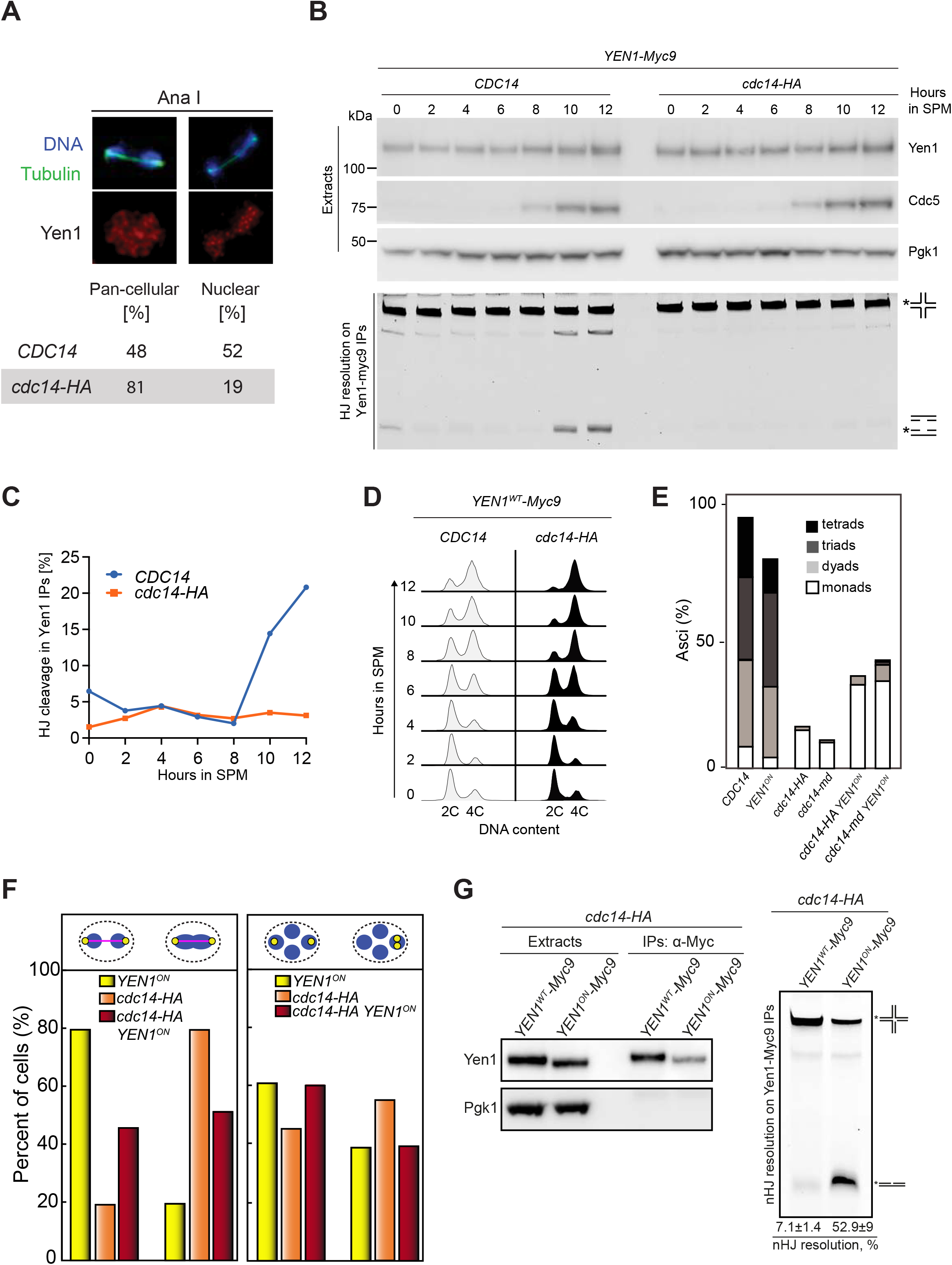
The absence of Cdc14 prevents activation of Yen1 during meiotic divisions. (A) Quantification of distinct localization patterns of Yen1 at meiosis I. (B) Analysis of expression levels and nuclease activity of Yen1 in *CDC14* (YJM7692) and *cdc14-HA* (YML7693) meiotic cells. Soluble extracts were prepared from *YEN1-Myc9* strains at 2-hr intervals after transfer into sporulation medium (SPM). Following anti-Myc immunoaffinity purification (IP), the IPs were analyzed by western blotting and tested for nuclease activity using Cy3-labeled Holliday junction DNA as a substrate. Upper panel: western blots of the cell extracts, with detection of Yen1-myc9, Cdc5, and Pgk1 (loading control). Lower panel: HJ resolution assay. The experiment shown is representative of two independent experiments. (C) Percentage of HJ cleavage in Yen1 IPs during meiosis in wild type and *cdc14-HA*. (D) Evolution of DNA content during meiosis from strains in (B). (E) Unrestrained resolution of recombination intermediates by Yen1^ON^ improves sporulation in *cdc14-HA* (JCY2164) and *cdc14-md* (PRY182) cells. Frequency of asci containing one, two, three and four spores in the strains of the indicated genotypes. (F) Frequency of cells presenting connected DAPI masses at anaphase I (left panel) or sister chromatid missegregation (right panel) in *YEN1^ON^* (PRY123/PRY99), *cdc14-HA* (JCY2327/PRY55) and *cdc14-HA YEN1^ON^* (PRY121/PRY108). (G) Poor HJ resolution in *cdc14-HA* (YML7693) is efficiently restored by the presence of Yen1^ON^ (JCY2421). Western blot analysis of Yen1/Yen1^ON^ immunoprecipitates is shown on the left panels. HJ resolution assay is shown on the right panel. Quantification of resolution efficiency is displayed at the bottom. Resolution data arise from two independent experiments.

### Constitutively active Yen1 supresses the accumulation of aberrant recombination intermediates in *cdc14* mutants

To determine whether the activation of Yen1 by Cdc14 is exerted by controlling its phosphorylation status, we next used a phosphorylation-resistant version of the nuclease (*YEN1^ON^*) in which mutation of nine of the CDK consensus sites of phosphorylation in Yen1 renders the protein constitutively active (Blanco et al. 2014; Arter et al. 2018). We combined the *cdc14-HA* allele with *YEN1^ON^* and examined sporulation efficiency. Similar to the elimination of Spo11-dependent DSBs, unrestrained activity of Yen1 partly rescued the sporulation defect of *cdc14-HA* (Fig 6E). We also directly tested the effect of *YEN1^ON^* in chromosome segregation during both meiotic divisions (Fig 6F). First, we checked if the presence of Yen1^ON^ promotes resolution of the DNA bridges observed at late anaphase I in *cdc14-HA* mutants. Indeed, a two-fold improvement in *cdc14-HA YEN1^ON^* over the *cdc14-HA* single mutant was observed (Fig 6F, left graph). Furthermore, missegregation of sister chromatids was also fully rescued to the levels of the *YEN1^ON^* single mutant in the *cdc14-HA YEN1^ON^* double mutant (Fig 6F, right graph). Moreover, to confirm if the improvement in chromosome segregation by *YEN1^ON^* was due to the reestablishment of Yen1 resolvase activity, we employed two types of assays. First, we checked using 2D-gels whether persistent accumulation of JMs and/or CO formation observed at the *HIS4LEU2* hotspot in the repair-deficient *sgs1-md mms4Δ cdc14-md* triple mutant was alleviated by the constitutively active Yen1^ON^ (Fig 5C-F). Notably, the introduction of *YEN1^ON^* efficiently prevented the accumulation of unresolved JMs after 24 hours in meiosis (Figs 5C and D). Furthermore, JMs were efficiently resolved giving rise to high levels of COs in the *sgs1-md mms4Δ cdc14-md YEN1^ON^* quadruple mutant (Figs 5E and F). Finally, we used the *in vitro* resolution assay to determine whether Yen1^ON^ was capable of restoring resolvase activity in the *cdc14-HA* mutant. We found that, unlike Yen1, the constitutively active Yen1^ON^ displayed potent resolvase activity in the absence of Cdc14 (Fig 6G). Altogether, these results indicate that Yen1 activation by Cdc14 is important for the repair of unprocessed recombination intermediates in meiosis.

### Cdc14 and Yen1 promote JM resolution during the first meiotic division

In the meiotic program, Yen1 activity appears to reach its maximum level during the second meiotic division (Matos et al. 2011; Arter et al. 2018). Nevertheless, it is possible that, during the first release of Cdc14 at meiosis I, Yen1 can be activated to act over remnants of unprocessed or complex JMs. In support of this idea we observed that 52% of wild-type cells displayed nuclear Yen1 in anaphase I (Fig 6A), indicating that a high proportion of Yen1 molecules have been translocated to the nucleus during the first meiotic division. To test if Cdc14 is acting over Yen1 already in meiosis I, or by contrast, Yen1 is exclusively activated during the second release of Cdc14 at meiosis II, we used the *cdc20-md* mutant to deplete the APC component Cdc20 and arrest cells in meiosis I (Carlile and Amon 2008). We analyzed the effect of Cdc14 and Yen1 on JM resolution in *cdc20-md* cells blocked at the first meiotic division when the Sgs1 and Mus81/Mms4 repair pathways, normally acting at the end of prophase I, have been removed. Remarkably, the *cdc20-md sgs1-md mms4Δ yen1Δ* and *cdc20-md sgs1-md mms4Δ cdc14-md* quadruple mutants exhibited much higher levels of unprocessed JMs than the *cdc20-md sgs1-md mms4Δ* triple mutant (Fig. 7A-B). In addition, this JM resolution defect was accompanied by a markedly reduced capacity of CO formation in both *cdc20-md sgs1-md mms4Δ yen1Δ* and *cdc20-md sgs1-md mms4Δ cdc14-md* quadruple mutants compared to the *cdc20-md sgs1-md mms4Δ* triple mutant (Fig. 7C-D). Altogether, we conclude that, when the action of other relevant JM processing pathways is compromised, Cdc14 and Yen1 are required for proper JM resolution as early as in the first meiotic division.

**Figure 7.**
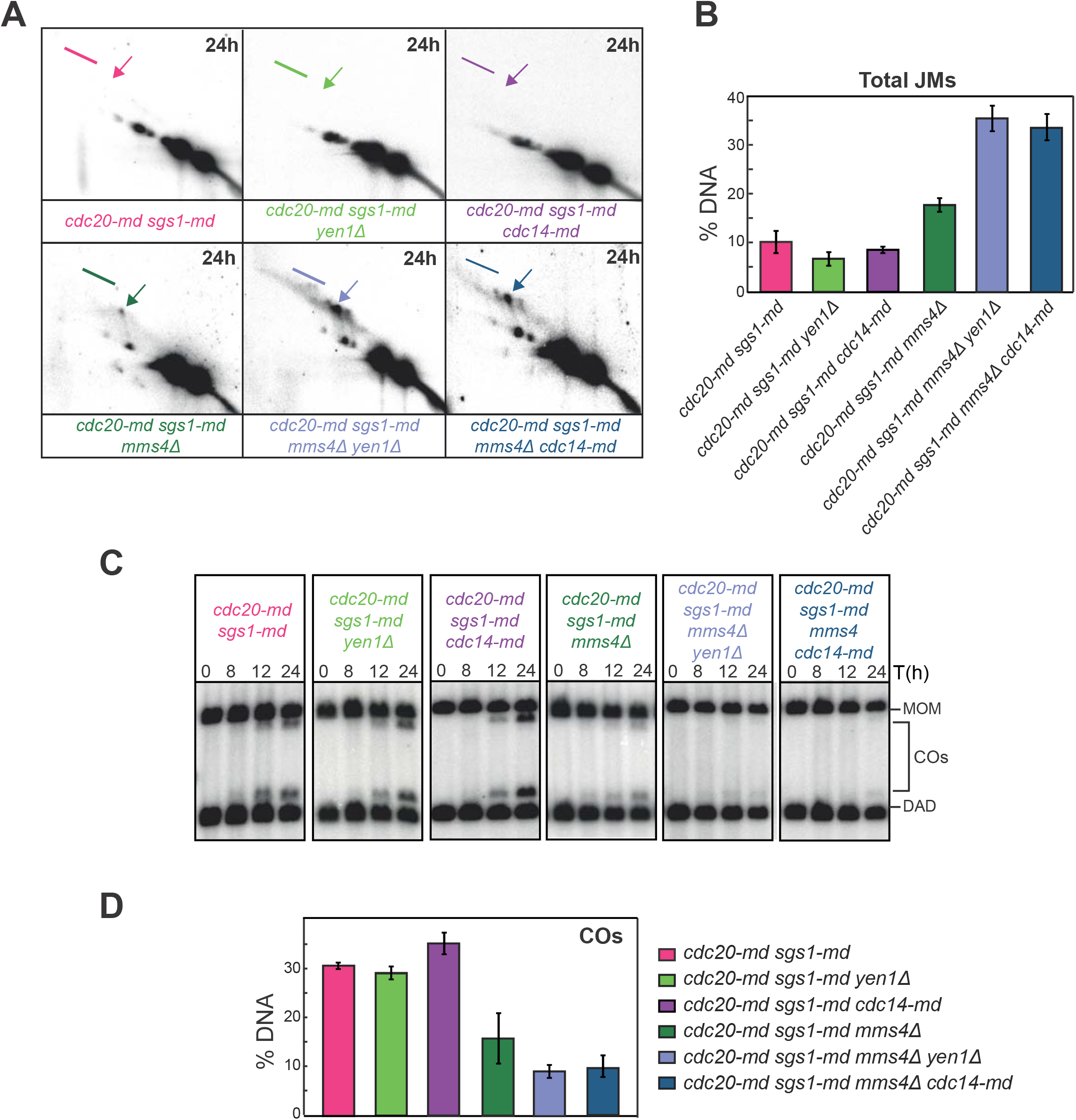
Analysis of recombination intermediates in the *cdc20-md* mutant reveals an early meiotic role for Cdc14 and Yen1. (A) Representative Southern blots depicting 2D-gels at the *HIS4LEU2* hotspot in *CDC14 cdc20-md sgs1-md* (JCY2480), *cdc20-md sgs1-md yen1* (JCY2469), *cdc20-md sgs1-md cdc14-md* (JCY2508), *cdc20-md sgs1-md mms4* (JCY2480), *cdc20-md sgs1-md mms4 yen1* (JCY2478) and *cdc20-md sgs1-md mms4 cdc14-md* (JCY2502). Arrows mark IH-dHJs and lines mark mc-JMs (see Fig 4A-B). (B) Quantification of total JMs in the strains shown in (A) from several independent gels. Error bars display SEM over the mean values plotted. (C) Representative 1D-gel Southern blot images for analysis of crossovers at the *HIS4LEU2* hotspot for all strains shown in (A). (D) Quantification of COs from at least three different image acquisitions like the one depicted in (C). Error bars represent SEM over the calculated mean value.

## Discussion

Meiotic recombination is fundamental for sexual reproduction to ensure that homologous chromosomes are accurately distributed to the gametes as well as to facilitate distinct allele combinations that sustain evolution. Homologous recombination is initiated by the introduction of programmed DSBs followed by resection, homology search, and DNA strand exchange. Those early steps in the recombination process lead to the formation of stable JMs, which are ultimately resolved into two main classes of HR repair products, known as COs and NCOs (Fig 8). Recombining chromosomes may also contain intermediates consisting of three-and four-armed DNA structures, such as mc-JMs, where three and four DNA duplexes stay connected (Schwacha and Kleckner 1995; Hunter and Kleckner 2001; Liberi et al. 2005; Cromie et al. 2006; Jessop and Lichten 2008; Oh et al. 2008; Bzymek et al. 2010; Mankouri et al. 2011). In some cases, unresolved recombination intermediates can persist until the metaphase to anaphase transition where a subset of late-acting nucleases take charge of their processing in order to safeguard genome integrity (San-Segundo and Clemente-Blanco 2020). Unrestrained activity of such nucleases can interfere with the correct allotment of COs and NCOs in meiosis (Arter et al. 2018); thus, they are tightly controlled by several layers of regulation (Fig. 8). Phosphorylation events carried out by key cell cycle kinases, like CDK, DDK, and Polo Kinase, are the most extended form of regulation of these enzymatic activities (Matos et al. 2011; Wild et al. 2019); therefore, it is expected that phosphatases can play also crucial roles in this regulation. In *S. cerevisiae* mitotic cells, at least one of the late acting nucleases, Yen1, is modulated by the highly-conserved Cdc14 phosphatase (Blanco et al. 2014; Eissler et al. 2014; Garcia-Luis et al. 2014). Although a number of studies have unveiled the relevance of Cdc14 during meiotic chromosome segregation and SPB/centrosome dynamics, a role for Cdc14 in the regulation of meiotic recombination has not been established so far. Given the essentiality of *CDC14* in the budding yeast, most previous meiotic studies have used conditional temperature-sensitive alleles of *cdc14* that are blocked in meiosis after the first round of chromosome segregation (Culotti and Hartwell 1971; Schild and Byers 1980; Marston et al. 2003; Fox et al. 2017). Despite the large amount of valuable data that has been collected over the years using these *ts* alleles, other less conspicuous functions of Cdc14, for example those affecting meiosis II, might have been precluded from being discovered. Here, using two different meiosis-specific alleles of *CDC14* we have been able to identify previously undetected functions of the phosphatase during meiosis, particularly those affecting meiotic recombination (Fig 8). Strikingly, we found the unprecedented requirement for Cdc14 to promote JM resolution at the transition between prophase I and the first meiotic division, suggesting that Cdc14 activity is not strictly ligated to its full release from the nucleolus.

**Figure 8.**
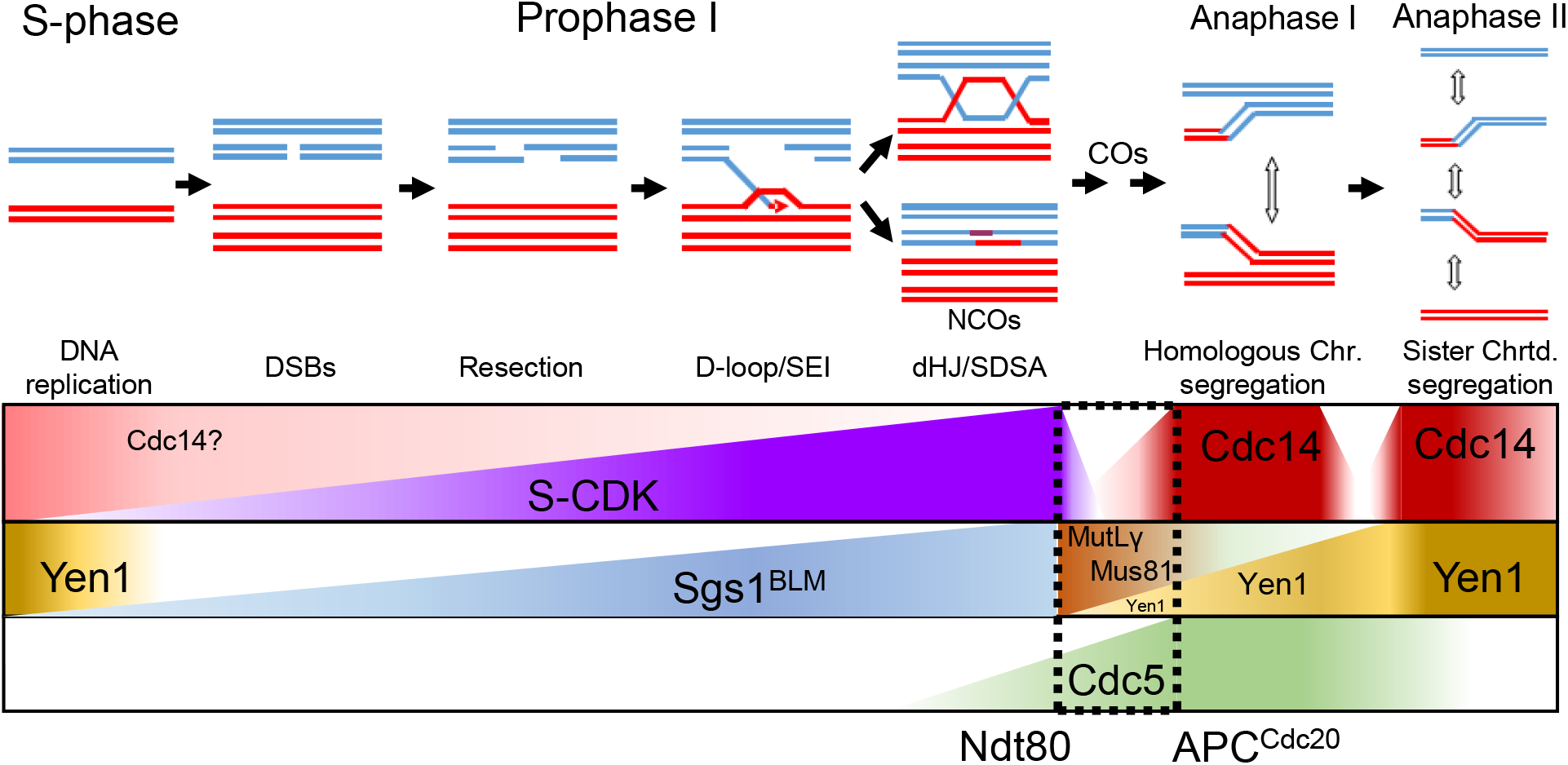
Model for *CDC14*-dependent resolution of recombination intermediates via multiple mechanisms. Contribution of Cdc14 to the correct disjunction of recombining chromosomes during meiosis. Early in prophase I, chromosomes initiate homologous recombination. 3’-resected ssDNA overhangs invade an intact template with the help of recombinases. Displacement of the intact strand from the template allows the formation of D-loops, which can be stabilized allowing DNA synthesis. Next, stabilized branched DNA molecules might be disrupted by the action of the anti-recombinases. Reannealing of the extended 3’-ssDNA overhang with the resected complementary strand, followed by further DNA synthesis will lead to the repair and ligation of the broken DNA duplex, giving rise to NCOs via SDSA. ZMM stabilization of JMs is followed by the resolution of dHJs through the MutLγ, class I, dedicated CO pathway. Ndt80-dependent expression, and activation, of Cdc5 triggers MutLγ resolvase activity. Unresolved linkages between bivalents that persist until metaphase/anaphase I are mostly resolved by the action of the SSEs, Mus81-Mms4. Slx1-Slx4, Top3-Rmi1 as well as Cdc14/Yen1 also contribute to the correct resolution of chromosomal entanglements between homologs during MI. Residual chromatid intertwining between sister chromatids during the second meiotic division will be removed by the action of Cdc14/Yen1. Gradual implementation of Yen1 activity during both divisions by Cdc14 will transfer Yen1 inactive population to its active/nuclear-enriched form.

### Novel insights into Cdc14 meiotic functions using different *cdc14* alleles

In the present work we describe an allele of *CDC14* (*cdc14-HA*) that displays no obvious defects during unchallenged mitotic divisions, but it is strongly deficient in sporulation. Protein levels of the phosphatase are reduced in both mitotic and meiotic *cdc14-HA* cells, but this reduction is more dramatic during meiosis (Fig 2A-D). However, the amount of the phosphatase in *cdc14-HA* meiotic cells is sufficient to complete meiotic DNA replication and both nuclear divisions with rather normal kinetics. Nonetheless, the *cdc14-HA* mutant fails to form spores once cells completed both nuclear divisions. It is possible that the impaired structural integrity of the SPBs observed in *cdc14-HA* can contribute to this phenotype. Similar defects in sporulation have been also described in the absence of components of the MEN pathway (Pablo-Hernando et al. 2007; Arguello-Miranda et al. 2017). Importantly, correct SPB re-duplication and separation occur in a high proportion of *cdc14-HA* cells, but not in the more restrictive *cdc14-md* mutant that contains negligible levels of Cdc14 during meiosis (Fig 2E-F; S3 Fig). Furthermore, many *cdc14-HA* cells are able to assemble functional tetrapolar spindles at metaphase/anaphase II confirming that problems arising in the *cdc14-HA* mutant somewhat differ from those occurring in *cdc14-md* as well as in the widely-employed, *cdc14-1, cdc14-3* or other FEAR mutants (Marston et al. 2003; Bizzari and Marston 2011; Fox et al. 2017). This observation suggests that the small amount of the phosphatase still present in *cdc14-HA* is sufficient to carry out those roles required for meiotic SPB dynamics, but not to maintain SPB integrity after divisions. The *cdc14-HA* mutant also displays a short delay in cohesin removal when entering anaphase I (Fig 3B-C and Fig S4D), sharing this particular meiotic phenotype with *spo12*, *slk19* and *cdc14-1* mutants (Marston et al. 2003). It is possible that delayed Rec8 removal from chromosomes could contribute to the presence of DNA bridges in late anaphase I. However, the proportion of *cdc14-HA* cells presenting anaphase I bridges is consistently higher than that of cells displaying persistent Rec8 signal during anaphase I; thus, it is unlikely that inefficient Rec8 removal is the only source of most DNA bridges, but instead a contributing factor. The fact that the formation of chromatin bridges in *cdc14-HA* and *cdc14-md* is substantially alleviated in the absence of meiotic DSBs strongly suggests that they may arise from chromosome entanglements resulting from unresolved recombination intermediates, and raises the possibility of a direct role for Cdc14 in the regulation of meiotic recombination that has not been previously explored. Here we demonstrate that, in some circumstances, Cdc14 is critically required for proper resolution of meiotic JMs through the regulation of the Yen1 nuclease.

### Multistage *CDC14*-dependent processing of recombination intermediates

Meiotic cells are endowed with a battery of enzymatic activities, including dissolvases and resolvases, that are tightly regulated to ensure efficient and timely processing of recombination intermediates (San-Segundo and Clemente-Blanco 2020). Processing of most meiotic JMs occurs at the end of pachytene (Borner et al. 2004), far earlier than the reported first release of Cdc14 at anaphase I. Nevertheless, this is not the first time that the requirement of Cdc14 has been linked to cell cycle stages preceding its bulk release from the nucleolus at anaphase. For example, DNA damage caused in vegetative cells triggers the transitory release of Cdc14 from the nucleolus to the nucleoplasm, allowing the phosphatase to act on components of the SPBs to stabilize them at metaphase (Villoria et al. 2017). Cdc14 is involved in completion of late-replicating regions in the rDNA, and other parts of the genome (Dulev et al. 2009). Although the *cdc14-HA* mutant does not show a discernible mitotic phenotype under unchallenged conditions, it is formally possible that *cdc14-HA* cells, which contain reduced protein levels, may accumulate aberrant DNA structures during preceding mitoses prior to meiosis entry. However, this is unlikely the case for the *cdc14-md* mutant, whose mitotic protein levels are very similar to those of the wild type. Therefore, we favor a role for Cdc14 in the direct regulation of substrates required for correct processing of meiotic JMs (Fig 8). Interestingly, the conserved SSE, Yen1^GEN1^ is a critical substrate of Cdc14 during budding yeast mitosis, and it exhibits a phosphorylation-regulated nucleocytoplasmic shuttling behaviour (Kosugi et al. 2009). CDK-dependent phosphorylation restricts Yen1 from entering the nucleus and becoming active, whereas reversal of that phosphorylation by Cdc14 allows Yen1 to enter the nucleus and resolve entangled DNA structures (Blanco et al. 2014; Eissler et al. 2014; Garcia-Luis et al. 2014). Nonetheless, Yen1 nuclear import and activation appears to be concomitant with bulk Cdc14 release from the nucleolus during anaphase. This is why a role for Yen1 in safeguarding chromosome segregation has been proposed, especially, during the second meiotic division (Matos et al. 2011; Blanco et al. 2014; Arter et al. 2018). Our results support this conclusion, since high frequencies of aneuploidy and missegregation events are detected during meiosis II in the *cdc14-HA* meiotic mutant when the other repair pathways are intact (Fig 3F). This is consistent with the requirement for Cdc14 to activate Yen1 (Fig 6A-C). Puzzlingly, we also observe Yen1-dependent processing of JMs in cells arrested during MI by Cdc20 depletion (Fig 7). This effect is manifested more prominently when the Mus81/Mms4 endonuclease and the Sgs1 helicase are missing. Such resolvase activity is not observed in *cdc14* mutants (Fig 7), suggesting that Cdc14 can activate Yen1 in cells that are about to initiate anaphase I. Taking into account that the massive Yen1 nuclear transport takes place during the ensuing anaphase (Arter et al. 2018), our results open the question of the origin of the Yen1 population acting in *CDC20*-depleted cells, which are stably arrested at the metaphase I to anaphase I transition in meiotic cells (Lee and Amon 2003; Katis et al. 2004; Carlile and Amon 2008; Arguello-Miranda et al. 2017). We speculate about other possible scenarios, like the existence of a different fraction of Yen1 arising from a different subcellular compartment to perform this earlier role. In support of this possibility, it has been recently postulated that a subpopulation of Yen1 localizes at the nucleolus in mitotic cells (Talhaoui et al. 2018). Whether the nuclear Yen1 observed in MI is sourced from the cytoplasm or from other sub-nuclear compartment will be a matter of future studies.

### Alternative repair pathways to process JMs during meiosis I

An intriguing possibility is that different SSEs might be required for processing different types of JMs that may not be necessarily good substrates for the canonical MutLγ complex. There is a risk that spindle forces originated during the *cdc20-md* arrest could generate tension at a number of still unprocessed JMs eventually forcing them towards conformations that cannot be easily processed by prophase-specific type of resolution/dissolution enzymes, for example affecting branch migration kinetics or directionality (Tang et al. 2015; Machin 2020). In such perilous scenario, Cdc14 could be an ideal regulator, together with Cdc5, to promote the switch between different endonucleases, by activating some while inhibiting others, once spindle forces start becoming dominant under circumstances where the SC no longer exists to hold homologs in close association (Shodhan et al. 2019; Cannavo et al. 2020; Kulkarni et al. 2020).

In budding yeast meiosis, the timing of JM resolution and CO formation is coordinated with cell-cycle progression through the *NDT80*-dependent expression of the Polo-like kinase Cdc5 (Clyne et al. 2003; Sourirajan and Lichten 2008; Matos et al. 2011). Thus, the requirement of Cdc5 for the resolution of dHJ at the pachytene to MI transition, upon Ndt80 activation, might too involve its ability to regulate the interaction of Cdc14 with Cif1/Net1 in the nucleolus. In opposition to mitotic cells, the action of Cdc5 could temporally counteract the negative regulatory effect of PP2A^Cdc55^ allowing some Cdc14 molecules to escape from its captor (Bizzari and Marston 2011; Kerr et al. 2011). It is tempting to speculate that Cdc5 might also play a relevant role during meiosis in promoting an early, metastable, partial release of a Cdc14 population at the transition from the pachytene stage to metaphase I in order to modulate the activity of a number of safeguarding enzymes required for correct chromosome segregation (Tsuchiya et al. 2014). Understanding the regulatory pathway/s which control Cdc14 activity previous to its full release during anaphase will be important to decipher its full contribution to the regulation of meiotic recombination.

In recent years, human orthologs of Cdc14 phosphatase have received increased attention due to their involvement in key processes like DDR, DNA repair and cell cycle control. Furthermore, recent findings point to recessive variants of the phosphatase to be directly responsible of human diseases, like deafness and male infertility (Imtiaz et al. 2018). Thus, in order to comprehend the underlying factors that trigger those conditions, a deeper understanding of the genetic and molecular mechanisms that are responsible for the countless functionalities of the Cdc14 phosphatase during gametogenesis and HR repair will be required.

## Materials and Methods

### Yeast strains and plasmids

All strains were SK1, as detailed in Table S1. Standard yeast manipulation procedures and growth media were utilized. To introduce the 3HA tag at the C-terminal end of Cdc14, the CDC14ha3-pKan^R^ plasmid containing the last ∼500bps of the *CDC14* gene with 3xHA inserted at the NotI restriction site and containing the *CLB1* terminator, was used. The plasmid was linearized using a unique restriction site located within the *CDC14* sequence and transformed into a SK1 haploid strain. Alternatively, *CDC14* was tagged using the PCR-based method described in (Longtine et al. 1998) using the plasmid pFA6a-3HA-kanMX6. The phenotype of *cdc14-HA* strains obtained from both tagging methods was checked and the sporulation defect was identical. Transformants containing correct tag integration were identified and tested by western blot for the presence of the tag. Southern blot analysis and/or PCR was performed to confirm the integration at the endogenous locus. Multi-copy plasmids used in Figure S2C were originally described in (Jaspersen et al. 1998).

### Synchronous meiotic time courses

Induction of synchronous meiosis was carried out according to the established protocols for standard assays (Padmore et al. 1991). All pre-growth and meiotic time courses were carried out at 30°C unless specified otherwise. For *cdc14-1* meiosis, the culture was kept at 23°C and shifted to 30°C 2 hours after transferring into sporulation medium (SPM). Aliquots were removed at the specified times and subjected to various analyses.

### DNA manipulation, extraction and southern blot detection

Standard DNA extraction was performed as in (Carballo et al. 2013). For studies at the *HIS4LEU2* recombination hotspot, the protocol described in (Ahuja and Borner 2011) was followed. For 2D gel agarose electrophoresis, cell cultures were photo-crosslinked with trioxalen (Merck) using long-wave UV light before DNA extraction, in order to visualize recombination intermediates by standard southern blotting techniques at the *HIS4LEU2* hotspot (Ahuja and Borner 2011).

### Time-lapse imaging, immunofluorescence, microscopy, and image analysis

Time-lapse experiments were performed as in (Newnham et al. 2013), with small variations. In brief, 1 ml aliquots from synchronous meiotic cultures were taken at specific times and diluted 1:9 in fresh SPM (kept at 30°C). 300μl of diluted cells were placed in suitable multi-well slides (80821 uncoated, ibidi). Slides were placed in a temperature-controlled incubation chamber from a Multidimensional Microscopy System Leica AF6000 LX. Images were taken at multiple positions and channels every 5, 10 or 15 minutes, depending on the experiment. Image acquisition was carried out using a CCD Hamamatsu 9100-02 camera. The system was controlled by the proprietary software, Leica Application Suite X, version 3.30. For preparations of fixed cells and immunofluorescence, aliquots were fixed and prepared as described in (Carballo et al. 2013) and (Arter et al. 2018), respectively. Chromosomal DNA was stained with 1μg/ml 4,6-diamino-2-phenylimide (DAPI). Images were recorded and analyzed using a Deltavision (DV3) workstation from Applied Precision Inc. with a Photometrics CoolSnap HQ (10-20MHz) air cooled CCD camera and controlled by SoftWorx image acquisition and deconvolution software.

### Protein extraction, Western blot analysis and antibodies

Whole-cell extracts were prepared from cell suspensions in 20% trichloroacetic acid by agitation with glass beads. Precipitated proteins were solubilized in SDS-PAGE sample buffer, and appropriate dilutions were analyzed by SDS-PAGE and western blotting (Gimenez-Abian et al. 2004). Antibodies used for western blotting were mouse monoclonal anti-MYC (1:1000, Abcam), mouse monoclonal anti-HA (1:1000) from S. Ley (NIMR), goat polyclonal anti-Cdc14 (yE-17; 1:1000; Santa Cruz Biotechnology), goat polyclonal anti-Cdc5 (1:1000; Santa Cruz Biotechnology), mouse monoclonal anti-Pgk1 (1:5000; Invitrogen), goat anti-mouse IgG conjugated to horseradish peroxidase (1:10000; Sigma-Aldrich), and chicken anti-rabbit IgG conjugated to horseradish peroxidase (1:10000; Sigma-Aldrich).

### Nuclease assays

For nuclease assays, myc9-tagged Yen1 and Yen1^ON^ were immuno-affinity purified from yeast using anti-Myc agarose beads (9E10) and washed extensively. The beads (approx. volume 10 μl) were then mixed with 10 μl cleavage buffer (50 mM Tris-HCl pH 7.5, 3 mM MgCl_2_) and 15 ng of 5’-Cy3-end-labeled synthetic Holliday junction X0 DNA. After 1 h incubation at 37°C with gentle rotation, reactions were stopped by addition of 2.5 μl of 10 mg/ml proteinase K and 2% SDS, followed by incubation for 30 min at 37°C. Loading buffer (3 μl) was then added and fluorescently-labelled products were separated by 10% native PAGE and analyzed using a Typhoon scanner and ImageQuant software. Resolution activity was calculated by determining the fraction of nicked duplex DNA product relative to the sum of the intact substrate and resolution product. The protein input was estimated by western blot.

### Data analysis and Biostatistics

Data was compiled and analyzed using Excel, LibreOffice Calc, and SPSS Statistical Data Editor. For multiple comparisons, analysis of variance (one-way ANOVA) was performed. For pairwise comparisons, two-tailed unpaired *t*-tests were used using IBM SPSS Statistics and SigmaPlot.

## Acknowledgements

We would like to thank C.R. Vázquez de Aldana, M. Knop, E. Winter, A.M. Neiman, E. Hoffmann, J.L. Santos, G.V. Börner, A. Marston, M.G. Blanco and F. Machín for kindly providing us with reagents, strains, antibodies and/or advice. We are especially thankful to the Molecular Biology of the Chromosome lab in CIB for sharing lab space, reagents and equipment. We would like to thank to the microscopy facility at the CIB for valuable technical advice in designing and analysing time-lapse, to P. Jalón for UV photography acquisition and to J.S. Ahuja, for advice in 2D gel experiments.

## Supplemental material legends

**Figure S1.**
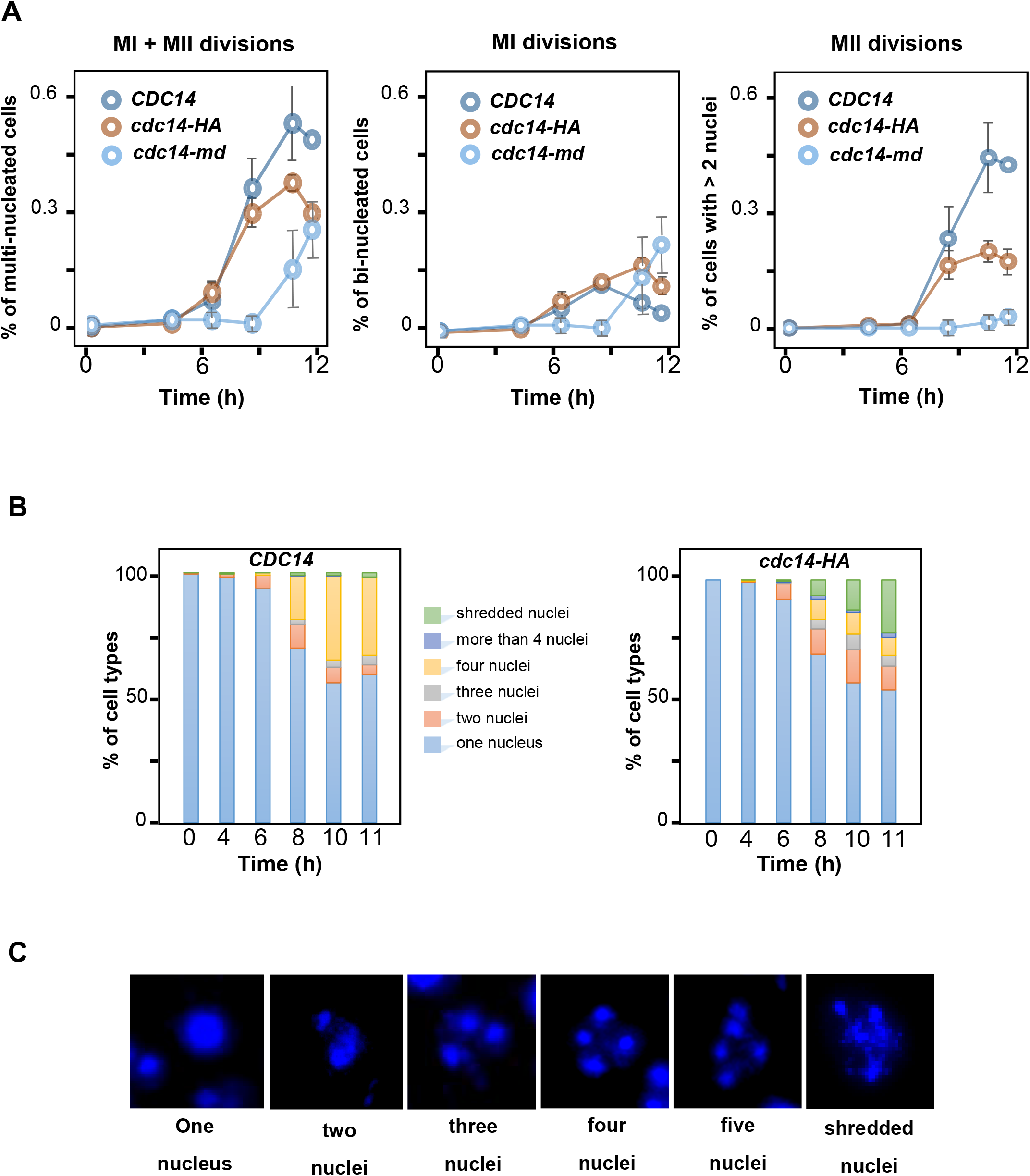
Cdc14 is required during meiosis for correct nuclear division. (A) *cdc14-HA* cells (JCY844) sustain both nuclear divisions although with subtly slower kinetics than wild type cells (JCY840) predominantly during the second round of nuclear segregation. *cdc14-md* cells (JCY2376) do not undergo the second nuclear division (B) Quantification of the number of nuclei as well as other aberrant nuclear structures during meiosis in *CDC14* (JCY840) and *cdc14-HA* (JCY844) cells. (C) Representative images of all nuclear structures quantified in (B).

**Figure S2.**
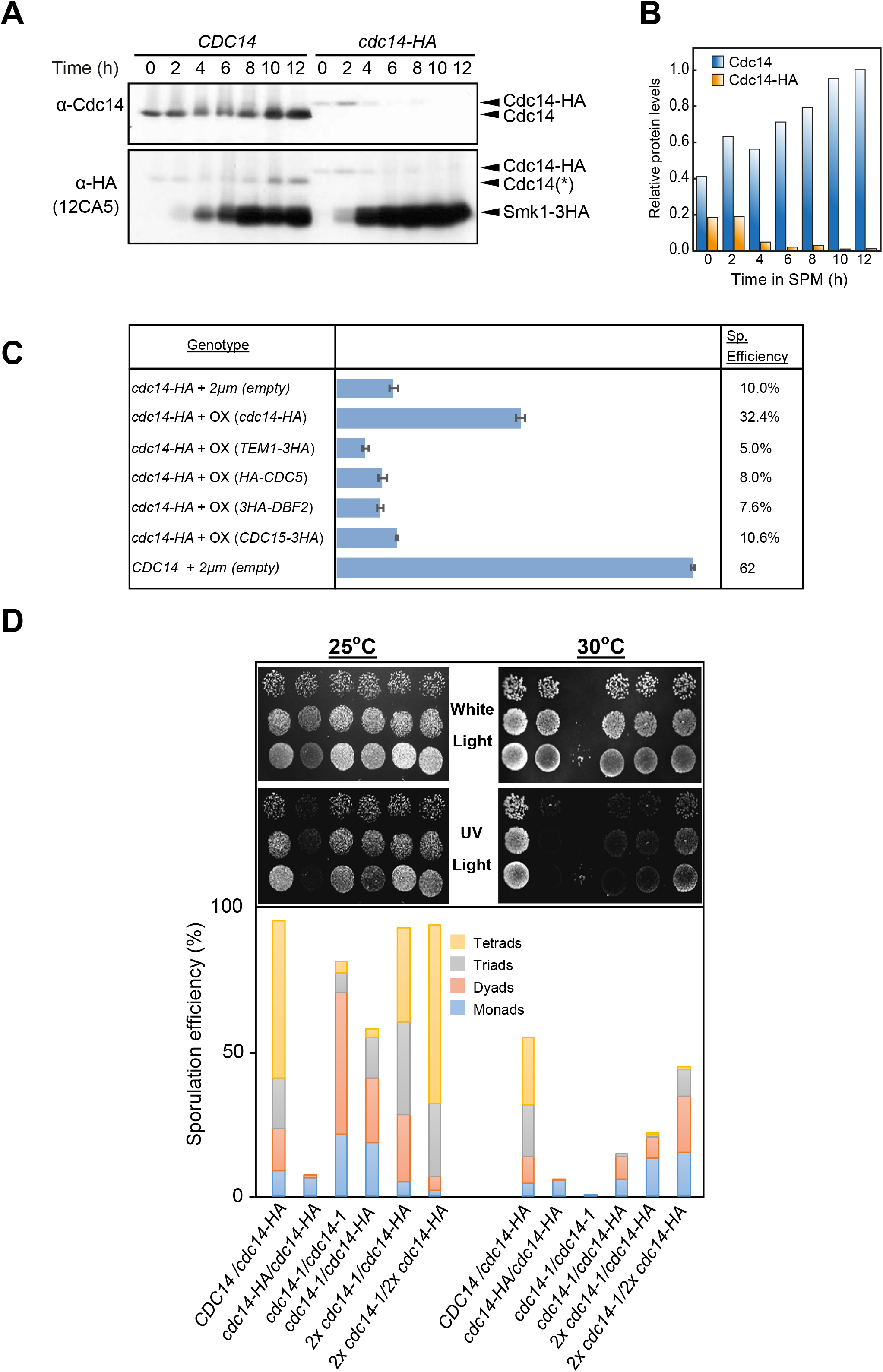
Reduced levels of Cdc14 causes the sporulation defect observed in the *cdc14-HA* mutant. (A) Analysis of Cdc14 protein levels in wild-type (JCY902), and *cdc14-HA* (JCY904) cells. Top panel shows immunodetection using a polyclonal α-Cdc14 (yE-17) antibody. The same blot was reanalyzed using α-HA (12CA5) (bottom panel). (*) denotes residual signal from the earlier α-Cdc14 detection. *SMK1-3HA* is also expressed in both *CDC14* and *cdc14-HA* strains. (B) Quantification of Cdc14 protein levels shown in (B). Values were normalized against the maximum signal corresponding to the 12 h time-point in the wild type (C) Sporulation efficiency of *cdc14-HA* diploids transformed with multi-copy plasmids carrying MEN genes (GGY102/GGY103/GGY104/GGY105) and *cdc14-HA* (GGY93). Only overproduction of the Cdc14 phosphatase rescues the sporulation defect. (D) Di-tyrosine autofluorescence of different mutant combinations as well as the control strains grown and sporulated on plates at 25°C and 30°C. *cdc14-1* homozygous diploids (JCY2353) sporulate at high efficiency under semi-permissive temperature forming preferentially tetrads whereas *cdc14-HA* homozygous diploids do not sporulate at any temperature (JCY840). Combinations, and variable copy number, of the mutant genes can rescue the sporulation defect at different degrees (JCY2365/JCY2354/JCY2356).

**Figure S3.**
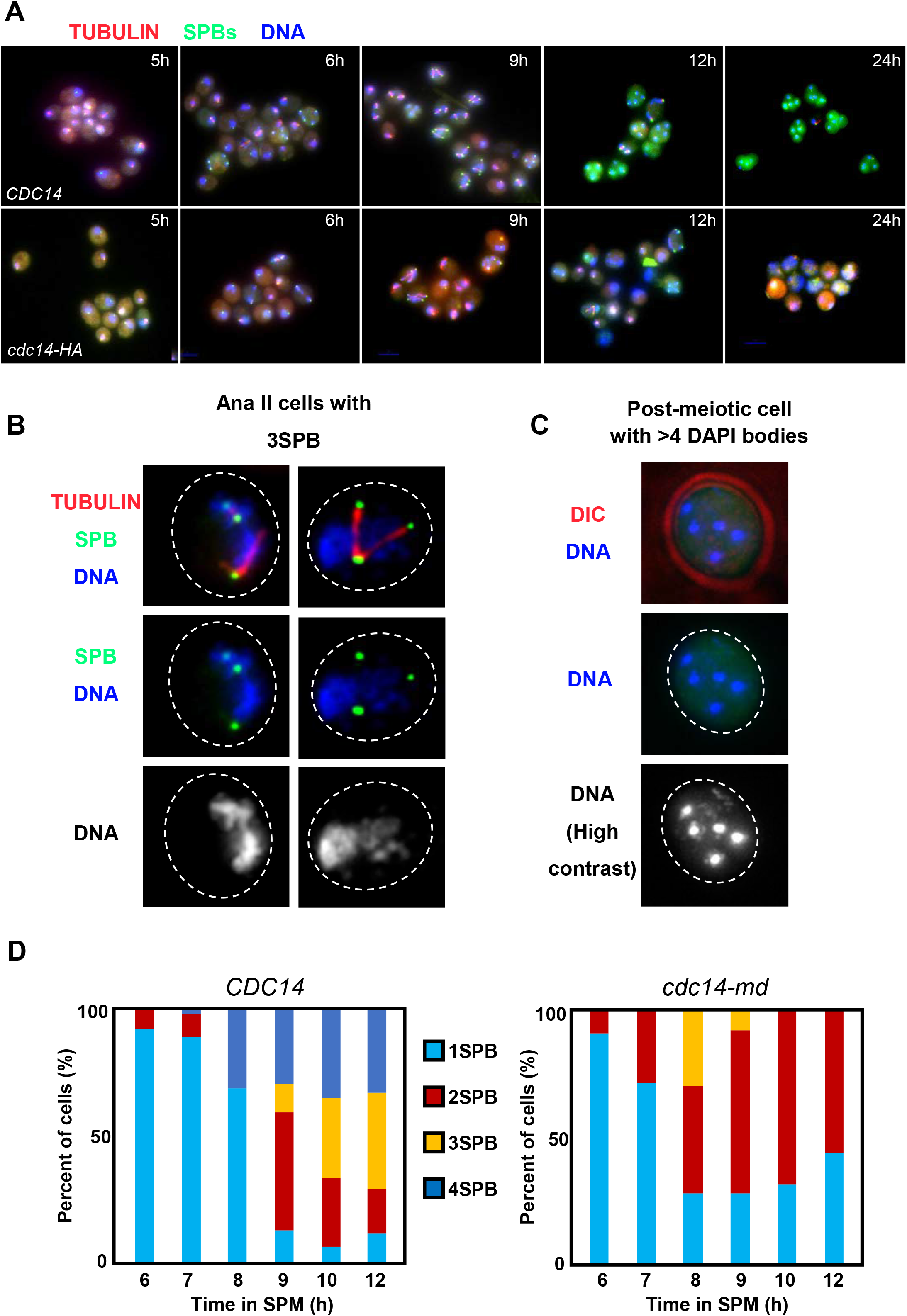
Meiotic *cdc14* mutant cells re-duplicate their SPBs and assemble meiosis I and meiosis II spindles. (A) Representative images of fixed cells at different time-points during synchronous parallel time courses for both *CDC14* (JCY892) and *cdc14* mutant (JCY893) cells. *cdc14-HA* cells initiate both meiotic divisions, as visualized by different fluorescence markers, but they do not complete sporulation. (B) Some *cdc14-HA* (JCY893) cells display abnormal number of SPBs and atypical spindle conformations. (C) Example of the terminal phenotype of a post-meiotic *cdc14-HA* (JCY893) cell. (D) *cdc14-md* cells display strong SPB re-duplication defects and arrest with two nuclei.

**Figure S4.**
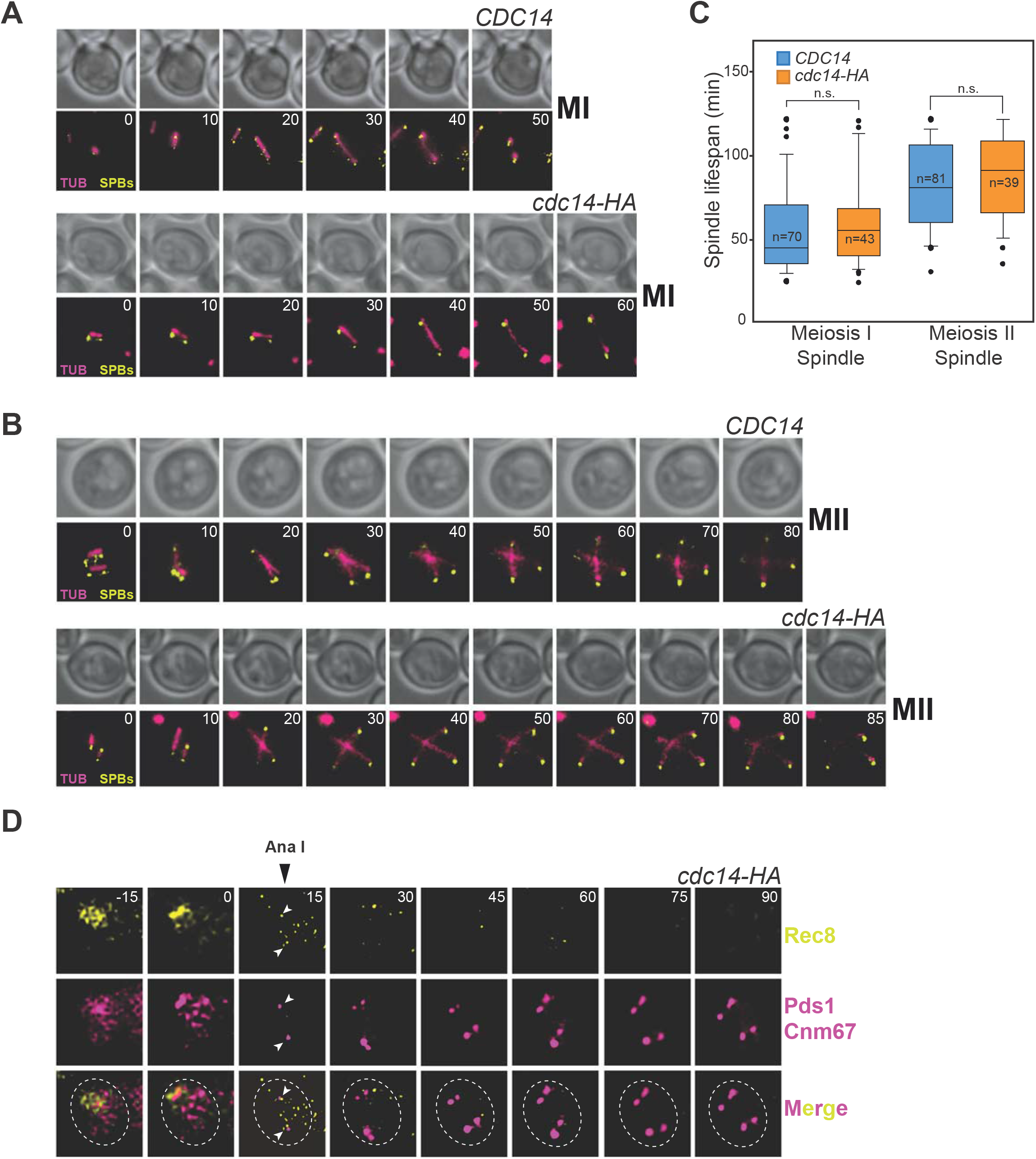
Meiotic spindle lifespan and dynamics are not altered in the meiosis-deficient *cdc14-HA* mutant. (A) Live-imaging of wild-type and *cdc14-HA* cells (GGY53 and GGY54, respectively) undergoing the first meiotic division carrying GFP-tubulin (red) and Spc29-CFP (green). (B) Visualization of spindle and SPB dynamics in the same strains (GGY53 and GGY54) undergoing the second meiotic division. For both A and B, microscopy fields from at least four different positions and from two separate wells were analyzed. A minimum of three time-courses per strain were run, acquired images were processed and movies generated (Video S1 and S2). Representative cells are shown. (C) Quantification of spindle lifespan from cells undergoing MI or MII. Meiotic deficient *cdc14-HA* cells (GGY54) show similar spindle dynamics compared to the wild type when completing both meiotic divisions. Box plots depict median number of spindle lifespan with whiskers representing upper and lower 1.5 interquartile range. Black dots represent outliers. Statistical test was performed using a two-tailed unpaired *t*-test. (D) Example of a meiotic *cdc14-HA* (JCY2404) cell transiting from prophase I to complete both meiotic divisions. Rec8-GFP (yellow) and Pds1-tdTomato/Cnm67-tdTomato (magenta). Time 0 (min) was considered the last frame where Pds1 was visualized and SPBs stayed in close association. Data obtained from Video S3 and S4.

**Figure S5.**
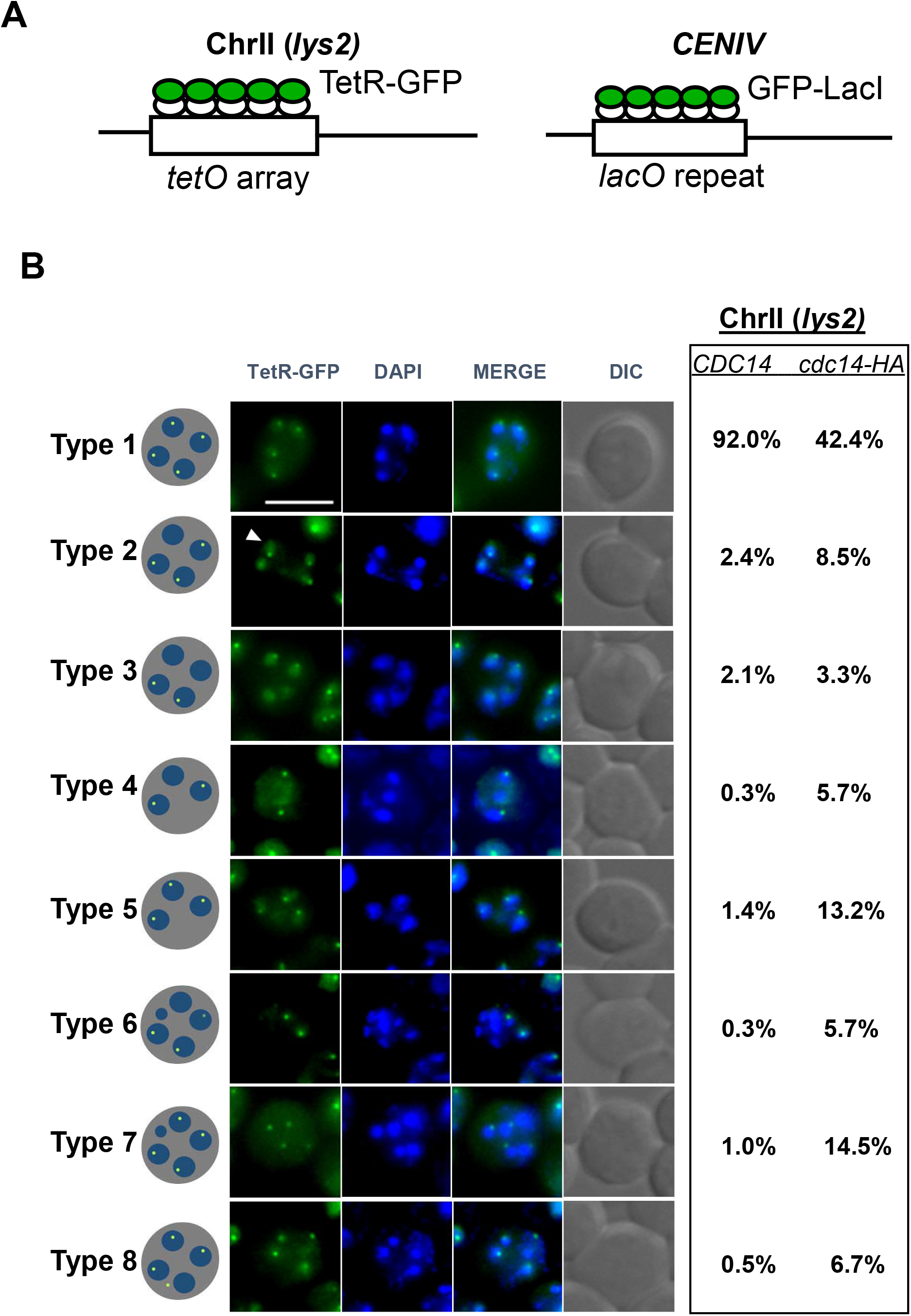
Lack of Cdc14 in meiosis causes missegregation of chromosomes at both meiotic divisions. (A) Scheme depicting the chromosome GFP-tagging system placed at an interstitial *locus* within chromosome II (left) and at a centromere proximal *locus* within chromosome IV (right). (B) Representative images and frequencies of the observed types of distribution of interstitial chromosome II GFP dots in homozygosis of both *cdc14-HA* mutant (JCY2230) and wild-type (JCY2231) meiotic cells. Arrowheads indicate diffuse GFP in the nucleus. For chromosome segregation fidelity analysis (Fig 3D), different percentages from depicted were calculated from considering only cells with four nuclei (type 1, 2 and 3). Cells were fixed in formaldehyde and nuclei stained with DAPI. Bar=5μm.

**Figure S6.**
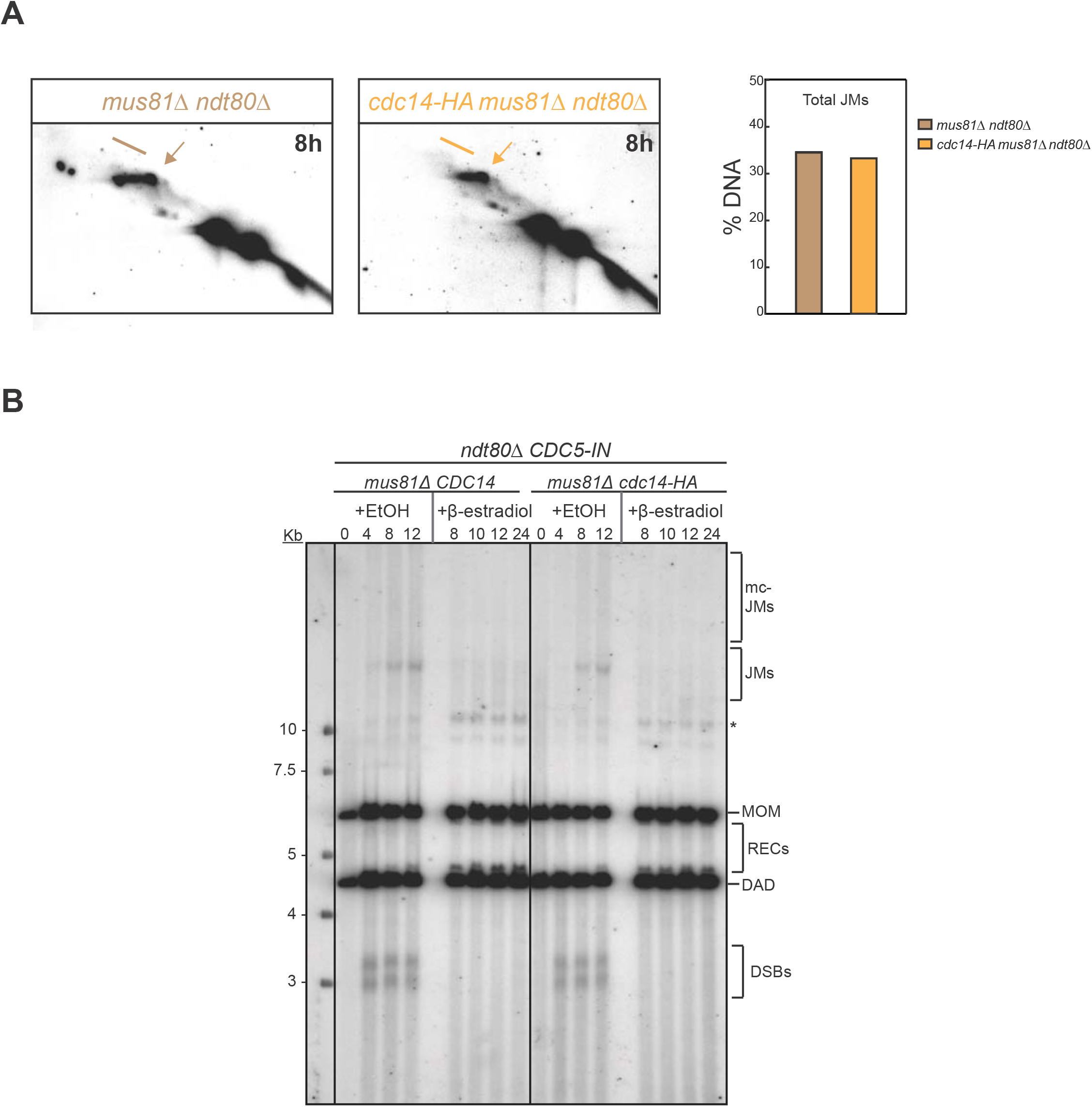
Accumulation of JMs is analogous in *cdc14-HA* in the presence or absence of Mus81. (A; left) Representative 2D gels showing recombination intermediates in *mus81 ndt80*, and *mus81 cdc14-HA ndt80* prophase-arrested cells 8 h after meiotic induction. (A; right) Quantification of DNA species from (2D-gels). (B) Comparison of JM abundance and resolution from the Southern blot shown in (Fig 5A). Asterisk indicates meiosis-specific non-characterized recombination products.

**Table S1.**
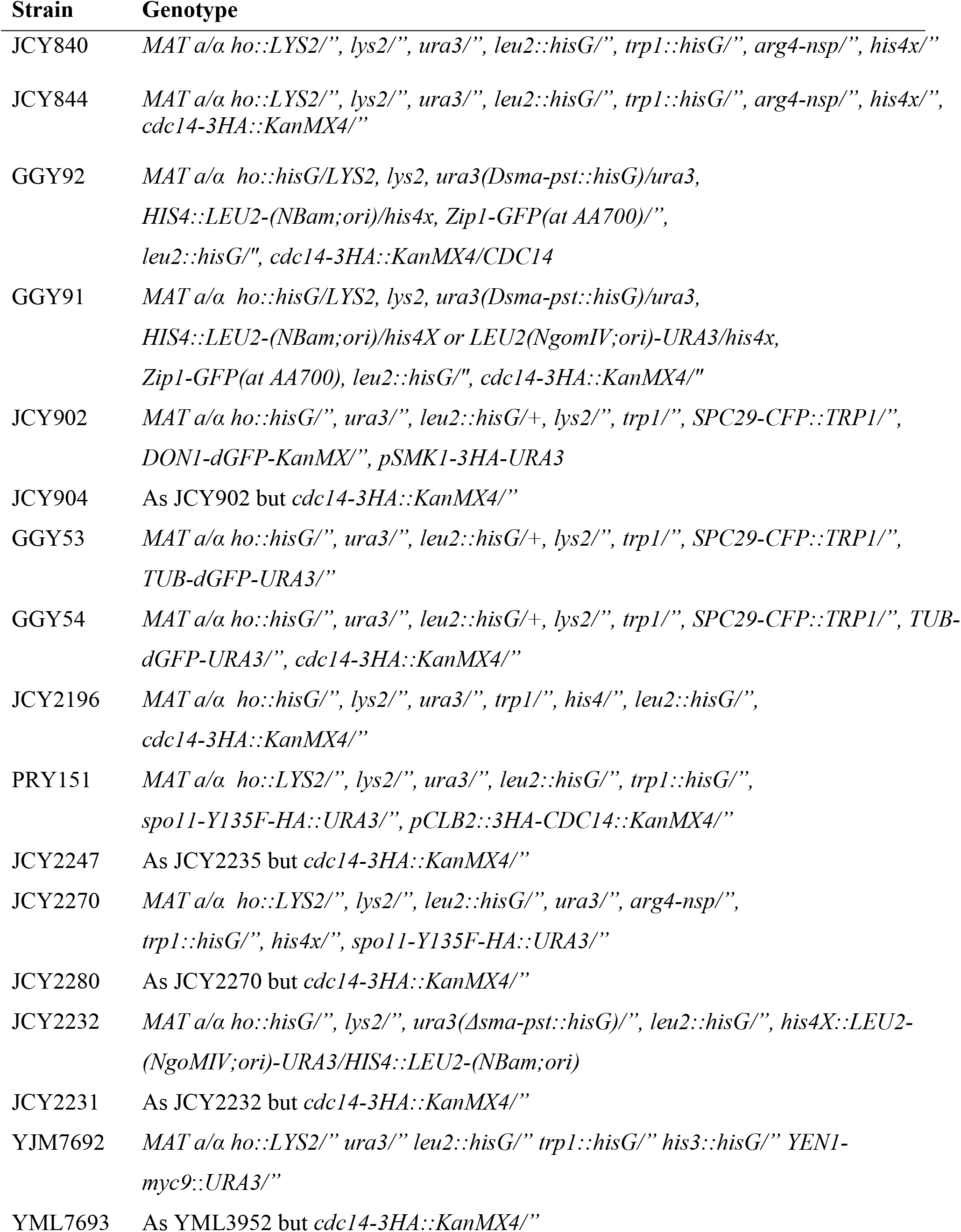
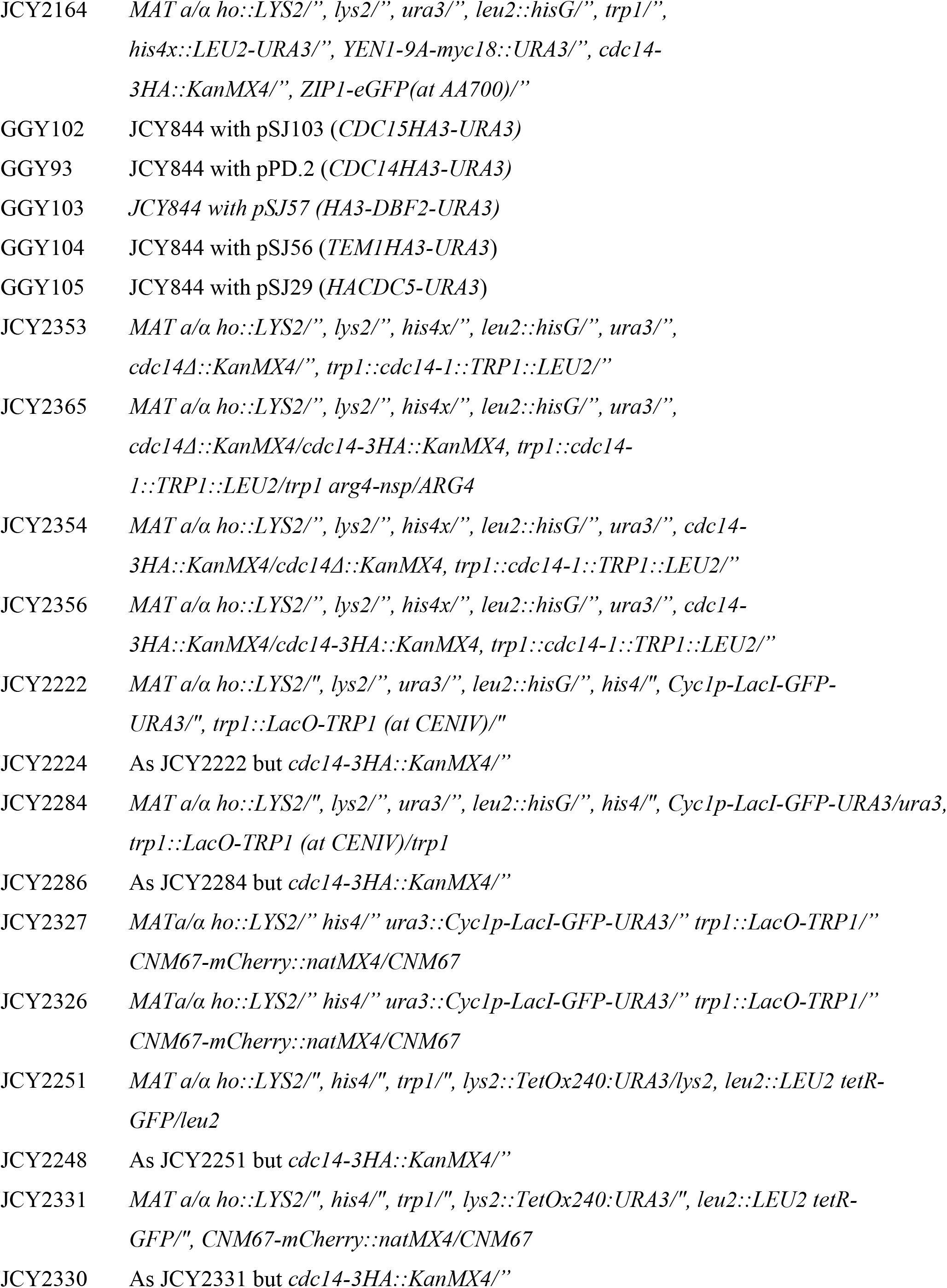
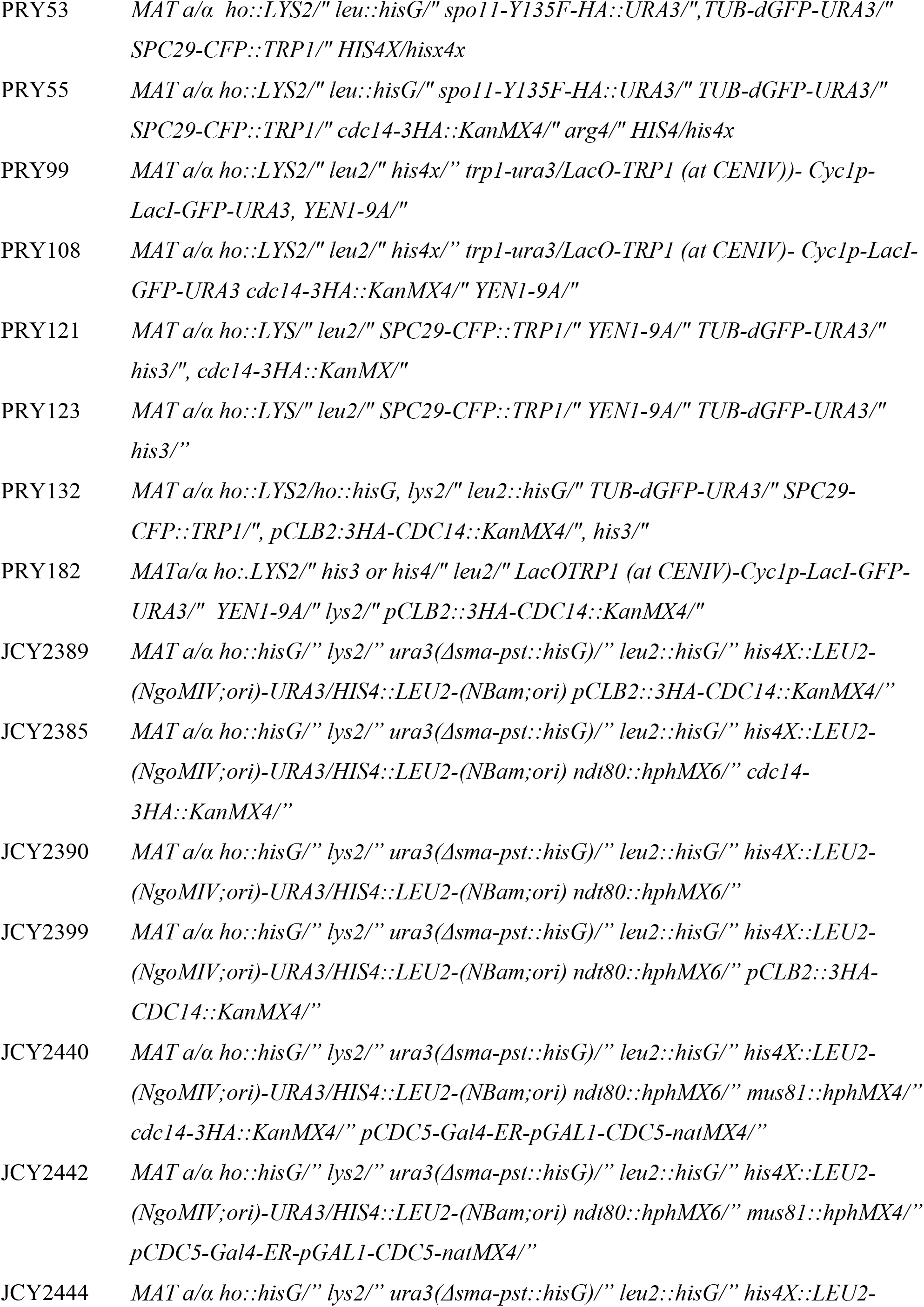
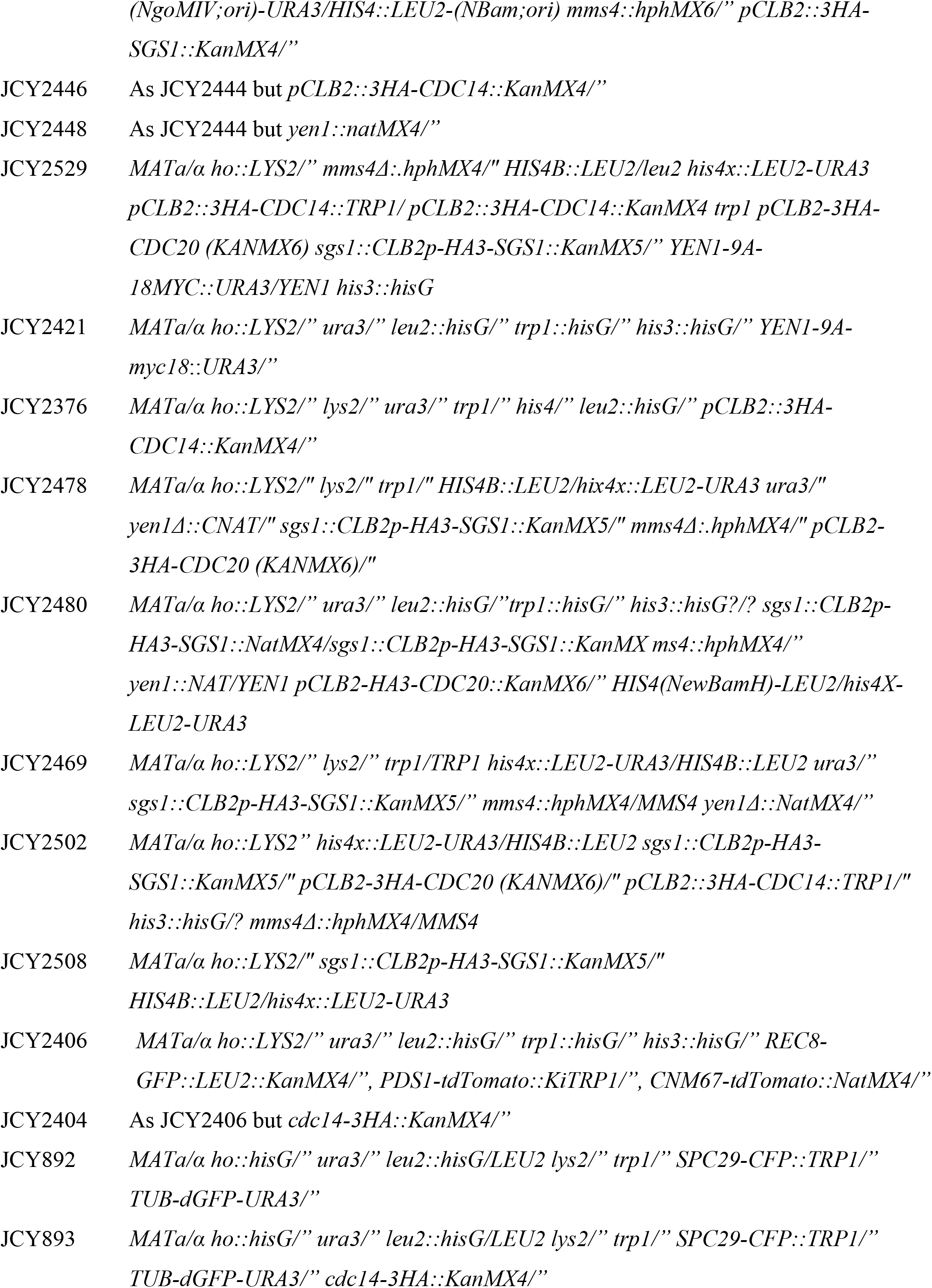
*S. cerevisiae* strains used in this study. SK1 background strains were used throughout the study, unless otherwise specified.

**Video S1. Meiotic spindle lifespan and dynamics in *CDC14* cells.** Live imaging of cells (GGY53) undergoing both meiotic divisions carrying GFP-tubulin (red) and Spc29-CFP (green).

**Video S2. Meiotic spindle lifespan and dynamics in *cdc14-HA* cells.** Live imaging of cells (GGY54) undergoing both meiotic divisions carrying GFP-tubulin (red) and Spc29-CFP (green).

**Video S3. Kinetics of Rec8-GFP association/removal in *CDC14* cells.** Sequence of Rec8-GFP wild-type cells (JCY2406) at 15-minute intervals displaying kinetics of cohesin assembly and removal.

**Video S4. Kinetics of Rec8-GFP association/removal in *cdc14-HA* cells.** Sequence of Rec8-GFP *cdc14-HA* cells (JCY2404) at 15 minutes interval displaying kinetics of cohesin removal.

## References

Ahuja JS, Borner GV. 2011. Analysis of meiotic recombination intermediates by two-dimensional gel electrophoresis. Methods Mol Biol 745: 99–116.

Allers T, Lichten M. 2001. Differential timing and control of noncrossover and crossover recombination during meiosis. Cell 106: 47–57.

Arguello-Miranda O, Zagoriy I, Mengoli V, Rojas J, Jonak K, Oz T, Graf P, Zachariae W. 2017. Casein Kinase 1 Coordinates Cohesin Cleavage, Gametogenesis, and Exit from M Phase in Meiosis II. Dev Cell 40: 37–52.

Arter M, Hurtado-Nieves V, Oke A, Zhuge T, Wettstein R, Fung JC, Blanco MG, Matos J. 2018. Regulated Crossing-Over Requires Inactivation of Yen1/GEN1 Resolvase during Meiotic Prophase I. Dev Cell 45: 785–800 e786.

Attner MA, Amon A. 2012. Control of the mitotic exit network during meiosis. Mol Biol Cell 23: 3122–3132.

Azzam R, Chen SL, Shou W, Mah AS, Alexandru G, Nasmyth K, Annan RS, Carr SA, Deshaies RJ. 2004. Phosphorylation by cyclin B-Cdk underlies release of mitotic exit activator Cdc14 from the nucleolus. Science 305: 516–519.

Bassermann F, Frescas D, Guardavaccaro D, Busino L, Peschiaroli A, Pagano M. 2008. The Cdc14B-Cdh1-Plk1 axis controls the G2 DNA-damage-response checkpoint. Cell 134: 256–267.

Bishop DK. 2006. Multiple mechanisms of meiotic recombination. Cell 127: 1095–1097.

Bizzari F, Marston AL. 2011. Cdc55 coordinates spindle assembly and chromosome disjunction during meiosis. J Cell Biol 193: 1213–1228.

Blanco MG, Matos J, West SC. 2014. Dual control of Yen1 nuclease activity and cellular localization by Cdk and Cdc14 prevents genome instability. Mol Cell 54: 94–106.

Bloom J, Cross FR. 2007. Novel role for Cdc14 sequestration: Cdc14 dephosphorylates factors that promote DNA replication. Mol Cell Biol 27: 842–853.

Borner GV, Kleckner N, Hunter N. 2004. Crossover/noncrossover differentiation, synaptonemal complex formation, and regulatory surveillance at the leptotene/zygotene transition of meiosis. Cell 117: 29–45.

Briza P, Winkler G, Kalchhauser H, Breitenbach M. 1986. Dityrosine is a prominent component of the yeast ascospore wall. A proof of its structure. J Biol Chem 261: 4288–4294.

Buonomo SB, Rabitsch KP, Fuchs J, Gruber S, Sullivan M, Uhlmann F, Petronczki M, Toth A, Nasmyth K. 2003. Division of the nucleolus and its release of CDC14 during anaphase of meiosis I depends on separase, SPO12, and SLK19. Dev Cell 4: 727–739.

Bzymek M, Thayer NH, Oh SD, Kleckner N, Hunter N. 2010. Double Holliday junctions are intermediates of DNA break repair. Nature 464: 937–941.

Cannavo E, Sanchez A, Anand R, Ranjha L, Hugener J, Adam C, Acharya A, Weyland N, Aran-Guiu X, Charbonnier JB et al. 2020. Regulation of the MLH1-MLH3 endonuclease in meiosis. Nature 586: 618–622.

Cao L, Alani E, Kleckner N. 1990. A pathway for generation and processing of double-strand breaks during meiotic recombination in S. cerevisiae. Cell 61: 1089–1101.

Carballo JA, Panizza S, Serrentino ME, Johnson AL, Geymonat M, Borde V, Klein F, Cha RS. 2013. Budding yeast ATM/ATR control meiotic double-strand break (DSB) levels by down-regulating Rec114, an essential component of the DSB-machinery. PLoS Genet 9: e1003545.

Carlile TM, Amon A. 2008. Meiosis I is established through division-specific translational control of a cyclin. Cell 133: 280–291.

Clemente-Blanco A, Mayan-Santos M, Schneider DA, Machin F, Jarmuz A, Tschochner H, Aragon L. 2009. Cdc14 inhibits transcription by RNA polymerase I during anaphase. Nature 458: 219–222.

Clemente-Blanco A, Sen N, Mayan-Santos M, Sacristan MP, Graham B, Jarmuz A, Giess A, Webb E, Game L, Eick D et al. 2011. Cdc14 phosphatase promotes segregation of telomeres through repression of RNA polymerase II transcription. Nat Cell Biol 13: 1450–1456.

Clyne RK, Katis VL, Jessop L, Benjamin KR, Herskowitz I, Lichten M, Nasmyth K. 2003. Polo-like kinase Cdc5 promotes chiasmata formation and cosegregation of sister centromeres at meiosis I. Nat Cell Biol 5: 480–485.

Copsey A, Tang S, Jordan PW, Blitzblau HG, Newcombe S, Chan AC, Newnham L, Li Z, Gray S, Herbert AD et al. 2013. Smc5/6 coordinates formation and resolution of joint molecules with chromosome morphology to ensure meiotic divisions. PLoS Genet 9: e1004071.

Cromie GA, Hyppa RW, Taylor AF, Zakharyevich K, Hunter N, Smith GR. 2006. Single Holliday junctions are intermediates of meiotic recombination. Cell 127: 1167–1178.

Culotti J, Hartwell LH. 1971. Genetic control of the cell division cycle in yeast. 3. Seven genes controlling nuclear division. Exp Cell Res 67: 389–401.

D’Amours D, Stegmeier F, Amon A. 2004. Cdc14 and condensin control the dissolution of cohesin-independent chromosome linkages at repeated DNA. Cell 117: 455–469.

de los Santos T, Loidl J, Larkin B, Hollingsworth NM. 2001. A role for MMS4 in the processing of recombination intermediates during meiosis in Saccharomyces cerevisiae. Genetics 159: 1511–1525.

De Muyt A, Jessop L, Kolar E, Sourirajan A, Chen J, Dayani Y, Lichten M. 2012. BLM helicase ortholog Sgs1 is a central regulator of meiotic recombination intermediate metabolism. Mol Cell 46: 43–53.

Diaz-Cuervo H, Bueno A. 2008. Cds1 controls the release of Cdc14-like phosphatase Flp1 from the nucleolus to drive full activation of the checkpoint response to replication stress in fission yeast. Mol Biol Cell 19: 2488–2499.

Dulev S, de Renty C, Mehta R, Minkov I, Schwob E, Strunnikov A. 2009. Essential global role of CDC14 in DNA synthesis revealed by chromosome underreplication unrecognized by checkpoints in cdc14 mutants. Proc Natl Acad Sci U S A 106: 14466–14471.

Eissler CL, Mazon G, Powers BL, Savinov SN, Symington LS, Hall MC. 2014. The Cdk/cDc14 module controls activation of the Yen1 holliday junction resolvase to promote genome stability. Mol Cell 54: 80–93.

Fox C, Zou J, Rappsilber J, Marston AL. 2017. Cdc14 phosphatase directs centrosome re-duplication at the meiosis I to meiosis II transition in budding yeast. Wellcome open research 2: 2.

Garcia-Luis J, Clemente-Blanco A, Aragon L, Machin F. 2014. Cdc14 targets the Holliday junction resolvase Yen1 to the nucleus in early anaphase. Cell Cycle 13: 1392–1399.

Gimenez-Abian MI, Rozalen AE, Carballo JA, Botella LM, Pincheira J, Lopez-Saez JF, de la Torre C. 2004. HSP90 and checkpoint-dependent lengthening of the G2 phase observed in plant cells under hypoxia and cold. Protoplasma 223: 191–196.

Grigaitis R, Ranjha L, Wild P, Kasaciunaite K, Ceppi I, Kissling V, Henggeler A, Susperregui A, Peter M, Seidel R et al. 2020. Phosphorylation of the RecQ Helicase Sgs1/BLM Controls Its DNA Unwinding Activity during Meiosis and Mitosis. Dev Cell 53: 706–723 e705.

Grigaitis R, Susperregui A, Wild P, Matos J. 2018. Characterization of DNA helicases and nucleases from meiotic extracts of S. cerevisiae. Methods Cell Biol 144: 371–388.

Hartwell LH, Mortimer RK, Culotti J, Culotti M. 1973. Genetic Control of the Cell Division Cycle in Yeast: V. Genetic Analysis of cdc Mutants. Genetics 74: 267–286.

Hartwell LH, Smith D. 1985. Altered fidelity of mitotic chromosome transmission in cell cycle mutants of S. cerevisiae. Genetics 110: 381–395.

Hunter N, Kleckner N. 2001. The single-end invasion: an asymmetric intermediate at the double-strand break to double-holliday junction transition of meiotic recombination. Cell 106: 59–70.

Imtiaz A, Belyantseva IA, Beirl AJ, Fenollar-Ferrer C, Bashir R, Bukhari I, Bouzid A, Shaukat U, Azaiez H, Booth KT et al. 2018. CDC14A phosphatase is essential for hearing and male fertility in mouse and human. Human molecular genetics 27: 780–798.

Ip SC, Rass U, Blanco MG, Flynn HR, Skehel JM, West SC. 2008. Identification of Holliday junction resolvases from humans and yeast. Nature 456: 357–361.

Jaspersen SL, Charles JF, Tinker-Kulberg RL, Morgan DO. 1998. A late mitotic regulatory network controlling cyclin destruction in Saccharomyces cerevisiae. Mol Biol Cell 9: 2803–2817.

Jessop L, Lichten M. 2008. Mus81/Mms4 endonuclease and Sgs1 helicase collaborate to ensure proper recombination intermediate metabolism during meiosis. Mol Cell 31: 313–323.

Kamieniecki RJ, Liu L, Dawson DS. 2005. FEAR but not MEN genes are required for exit from meiosis I. Cell Cycle 4: 1093–1098.

Kamieniecki RJ, Shanks RM, Dawson DS. 2000. Slk19p is necessary to prevent separation of sister chromatids in meiosis I. Curr Biol 10: 1182–1190.

Katis VL, Lipp JJ, Imre R, Bogdanova A, Okaz E, Habermann B, Mechtler K, Nasmyth K, Zachariae W. 2010. Rec8 phosphorylation by casein kinase 1 and Cdc7-Dbf4 kinase regulates cohesin cleavage by separase during meiosis. Dev Cell 18: 397–409.

Katis VL, Matos J, Mori S, Shirahige K, Zachariae W, Nasmyth K. 2004. Spo13 facilitates monopolin recruitment to kinetochores and regulates maintenance of centromeric cohesion during yeast meiosis. Curr Biol 14: 2183–2196.

Keeney S, Giroux CN, Kleckner N. 1997. Meiosis-specific DNA double-strand breaks are catalyzed by Spo11, a member of a widely conserved protein family. Cell 88: 375–384.

Kerr GW, Sarkar S, Tibbles KL, Petronczki M, Millar JB, Arumugam P. 2011. Meiotic nuclear divisions in budding yeast require PP2A(Cdc55)-mediated antagonism of Net1 phosphorylation by Cdk. J Cell Biol 193: 1157–1166.

Kosugi S, Hasebe M, Tomita M, Yanagawa H. 2009. Systematic identification of cell cycle-dependent yeast nucleocytoplasmic shuttling proteins by prediction of composite motifs. Proc Natl Acad Sci U S A 106: 10171–10176.

Kulkarni DS, Owens SN, Honda M, Ito M, Yang Y, Corrigan MW, Chen L, Quan AL, Hunter N. 2020. PCNA activates the MutLgamma endonuclease to promote meiotic crossing over. Nature 586: 623–627.

Lee BH, Amon A. 2003. Role of Polo-like kinase CDC5 in programming meiosis I chromosome segregation. Science 300: 482–486.

Li L, Ernsting BR, Wishart MJ, Lohse DL, Dixon JE. 1997. A family of putative tumor suppressors is structurally and functionally conserved in humans and yeast. J Biol Chem 272: 29403–29406.

Liberi G, Maffioletti G, Lucca C, Chiolo I, Baryshnikova A, Cotta-Ramusino C, Lopes M, Pellicioli A, Haber JE, Foiani M. 2005. Rad51-dependent DNA structures accumulate at damaged replication forks in sgs1 mutants defective in the yeast ortholog of BLM RecQ helicase. Genes Dev 19: 339–350.

Longtine MS, McKenzie A, 3rd, Demarini DJ, Shah NG, Wach A, Brachat A, Philippsen P, Pringle JR. 1998. Additional modules for versatile and economical PCR-based gene deletion and modification in Saccharomyces cerevisiae. Yeast 14: 953–961.

Machin F. 2020. Implications of Metastable Nicks and Nicked Holliday Junctions in Processing Joint Molecules in Mitosis and Meiosis. Genes 11.

Machin F, Torres-Rosell J, De Piccoli G, Carballo JA, Cha RS, Jarmuz A, Aragon L. 2006. Transcription of ribosomal genes can cause nondisjunction. J Cell Biol 173: 893–903.

Mankouri HW, Ashton TM, Hickson ID. 2011. Holliday junction-containing DNA structures persist in cells lacking Sgs1 or Top3 following exposure to DNA damage. Proc Natl Acad Sci U S A 108: 4944–4949.

Marston AL, Lee BH, Amon A. 2003. The Cdc14 phosphatase and the FEAR network control meiotic spindle disassembly and chromosome segregation. Dev Cell 4: 711–726.

Martini E, Diaz RL, Hunter N, Keeney S. 2006. Crossover homeostasis in yeast meiosis. Cell 126: 285–295.

Matos J, Blanco MG, Maslen S, Skehel JM, West SC. 2011. Regulatory control of the resolution of DNA recombination intermediates during meiosis and mitosis. Cell 147: 158–172.

Michaelis C, Ciosk R, Nasmyth K. 1997. Cohesins: Chromosomal Proteins that Prevent Premature Separation of Sister Chromatids. Cell 91: 35–45.

Mocciaro A, Berdougo E, Zeng K, Black E, Vagnarelli P, Earnshaw W, Gillespie D, Jallepalli P, Schiebel E. 2010. Vertebrate cells genetically deficient for Cdc14A or Cdc14B retain DNA damage checkpoint proficiency but are impaired in DNA repair. J Cell Biol 189: 631–639.

Mocciaro A, Schiebel E. 2010. Cdc14: a highly conserved family of phosphatases with non-conserved functions? J Cell Sci 123: 2867–2876.

Newnham L, Jordan PW, Carballo JA, Newcombe S, Hoffmann E. 2013. Ipl1/Aurora kinase suppresses S-CDK-driven spindle formation during prophase I to ensure chromosome integrity during meiosis. PLoS One 8: e83982.

Nolt JK, Rice LM, Gallo-Ebert C, Bisher ME, Nickels JT. 2011. PP2A (Cdc)(5)(5) is required for multiple events during meiosis I. Cell Cycle 10: 1420–1434.

Oh SD, Lao JP, Hwang PY, Taylor AF, Smith GR, Hunter N. 2007. BLM ortholog, Sgs1, prevents aberrant crossing-over by suppressing formation of multichromatid joint molecules. Cell 130: 259–272.

Oh SD, Lao JP, Taylor AF, Smith GR, Hunter N. 2008. RecQ helicase, Sgs1, and XPF family endonuclease, Mus81-Mms4, resolve aberrant joint molecules during meiotic recombination. Mol Cell 31: 324–336.

Pablo-Hernando ME, Arnaiz-Pita Y, Nakanishi H, Dawson D, del Rey F, Neiman AM, Vazquez de Aldana CR. 2007. Cdc15 is required for spore morphogenesis independently of Cdc14 in Saccharomyces cerevisiae. Genetics 177: 281–293.

Padmore R, Cao L, Kleckner N. 1991. Temporal comparison of recombination and synaptonemal complex formation during meiosis in S. cerevisiae. Cell 66: 1239–1256.

Piazza A, Wright WD, Heyer WD. 2017. Multi-invasions Are Recombination Byproducts that Induce Chromosomal Rearrangements. Cell 170: 760–773 e715.

Queralt E, Lehane C, Novak B, Uhlmann F. 2006. Downregulation of PP2A(Cdc55) phosphatase by separase initiates mitotic exit in budding yeast. Cell 125: 719–732.

Rahal R, Amon A. 2008. The Polo-like kinase Cdc5 interacts with FEAR network components and Cdc14. Cell Cycle 7: 3262–3272.

Rodriguez-Rodriguez JA, Moyano Y, Jativa S, Queralt E. 2016. Mitotic Exit Function of Polo-like Kinase Cdc5 Is Dependent on Sequential Activation by Cdk1. Cell reports 15: 2050–2062.

Rosso L, Marques AC, Weier M, Lambert N, Lambot MA, Vanderhaeghen P, Kaessmann H. 2008. Birth and rapid subcellular adaptation of a hominoid-specific CDC14 protein. PLoS Biol 6: e140.

San-Segundo PA, Clemente-Blanco A. 2020. Resolvases, Dissolvases, and Helicases in Homologous Recombination: Clearing the Road for Chromosome Segregation. Genes 11.

Sanchez A, Adam C, Rauh F, Duroc Y, Ranjha L, Lombard B, Mu X, Wintrebert M, Loew D, Guarne A et al. 2020. Exo1 recruits Cdc5 polo kinase to MutLgamma to ensure efficient meiotic crossover formation. Proc Natl Acad Sci U S A 117: 30577–30588.

Schild D, Byers B. 1980. Diploid spore formation and other meiotic effects of two cell-division-cycle mutations of Saccharomyces cerevisiae. Genetics 96: 859–876.

Schwacha A, Kleckner N. 1994. Identification of joint molecules that form frequently between homologs but rarely between sister chromatids during yeast meiosis. Cell 76: 51–63.

Schwacha A, Kleckner N. 1995. Identification of double Holliday junctions as intermediates in meiotic recombination. Cell 83: 783–791.

Sharon G, Simchen G. 1990. Mixed segregation of chromosomes during single-division meiosis of Saccharomyces cerevisiae. Genetics 125: 475–485.

Shinohara M, Oh SD, Hunter N, Shinohara A. 2008. Crossover assurance and crossover interference are distinctly regulated by the ZMM proteins during yeast meiosis. Nat Genet 40: 299–309.

Shodhan A, Medhi D, Lichten M. 2019. Noncanonical Contributions of MutLgamma to VDE-Initiated Crossovers During Saccharomyces cerevisiae Meiosis. G3 (Bethesda, Md) 9: 1647–1654.

Shonn MA, McCarroll R, Murray AW. 2002. Spo13 protects meiotic cohesin at centromeres in meiosis I. Genes Dev 16: 1659–1671.

Shou W, Seol JH, Shevchenko A, Baskerville C, Moazed D, Chen ZW, Jang J, Shevchenko A, Charbonneau H, Deshaies RJ. 1999. Exit from mitosis is triggered by Tem1-dependent release of the protein phosphatase Cdc14 from nucleolar RENT complex. Cell 97: 233–244.

Sourirajan A, Lichten M. 2008. Polo-like kinase Cdc5 drives exit from pachytene during budding yeast meiosis. Genes Dev 22: 2627–2632.

Stegmeier F, Visintin R, Amon A. 2002. Separase, polo kinase, the kinetochore protein Slk19, and Spo12 function in a network that controls Cdc14 localization during early anaphase. Cell 108: 207–220.

Straight AF, Belmont AS, Robinett CC, Murray AW. 1996. GFP tagging of budding yeast chromosomes reveals that protein-protein interactions can mediate sister chromatid cohesion. Curr Biol 6: 1599–1608.

Sullivan M, Higuchi T, Katis VL, Uhlmann F. 2004. Cdc14 phosphatase induces rDNA condensation and resolves cohesin-independent cohesion during budding yeast anaphase. Cell 117: 471–482.

Talhaoui I, Bernal M, Mullen JR, Dorison H, Palancade B, Brill SJ, Mazon G. 2018. Slx5-Slx8 ubiquitin ligase targets active pools of the Yen1 nuclease to limit crossover formation. Nat Commun 9: 5016.

Tang S, Wu MKY, Zhang R, Hunter N. 2015. Pervasive and essential roles of the Top3-Rmi1 decatenase orchestrate recombination and facilitate chromosome segregation in meiosis. Mol Cell 57: 607–621.

Torres-Rosell J, Machin F, Jarmuz A, Aragon L. 2004. Nucleolar segregation lags behind the rest of the genome and requires Cdc14p activation by the FEAR network. Cell Cycle 3: 496–502.

Trautmann S, McCollum D. 2002. Cell cycle: new functions for Cdc14 family phosphatases. Curr Biol 12: R733–735.

Tsuchiya D, Yang Y, Lacefield S. 2014. Positive feedback of NDT80 expression ensures irreversible meiotic commitment in budding yeast. PLoS Genet 10: e1004398.

Villoria MT, Ramos F, Duenas E, Faull P, Cutillas PR, Clemente-Blanco A. 2017. Stabilization of the metaphase spindle by Cdc14 is required for recombinational DNA repair. EMBO J 36: 79–101.

Visintin C, Tomson BN, Rahal R, Paulson J, Cohen M, Taunton J, Amon A, Visintin R. 2008. APC/C-Cdh1-mediated degradation of the Polo kinase Cdc5 promotes the return of Cdc14 into the nucleolus. Genes Dev 22: 79–90.

Visintin R, Craig K, Hwang ES, Prinz S, Tyers M, Amon A. 1998. The phosphatase Cdc14 triggers mitotic exit by reversal of Cdk-dependent phosphorylation. Mol Cell 2: 709–718.

Visintin R, Hwang ES, Amon A. 1999. Cfi1 prevents premature exit from mitosis by anchoring Cdc14 phosphatase in the nucleolus. Nature 398: 818–823.

Visintin R, Stegmeier F, Amon A. 2003. The role of the polo kinase Cdc5 in controlling Cdc14 localization. Mol Biol Cell 14: 4486–4498.

Wang BD, Yong-Gonzalez V, Strunnikov AV. 2004. Cdc14p/FEAR pathway controls segregation of nucleolus in S. cerevisiae by facilitating condensin targeting to rDNA chromatin in anaphase. Cell Cycle 3: 960–967.

Wild P, Susperregui A, Piazza I, Dorig C, Oke A, Arter M, Yamaguchi M, Hilditch AT, Vuina K, Chan KC et al. 2019. Network Rewiring of Homologous Recombination Enzymes during Mitotic Proliferation and Meiosis. Mol Cell 75: 859–874 e854.

Xaver M, Huang L, Chen D, Klein F. 2013. Smc5/6-Mms21 prevents and eliminates inappropriate recombination intermediates in meiosis. PLoS Genet 9: e1004067.

Yellman CM, Roeder GS. 2015. Cdc14 Early Anaphase Release, FEAR, Is Limited to the Nucleus and Dispensable for Efficient Mitotic Exit. PLoS One 10: e0128604.

Yoshida S, Toh-e A. 2002. Budding yeast Cdc5 phosphorylates Net1 and assists Cdc14 release from the nucleolus. Biochem Biophys Res Commun 294: 687–691.

Zakharyevich K, Ma Y, Tang S, Hwang PY, Boiteux S, Hunter N. 2010. Temporally and biochemically distinct activities of Exo1 during meiosis: double-strand break resection and resolution of double Holliday junctions. Mol Cell 40: 1001–1015.

Zakharyevich K, Tang S, Ma Y, Hunter N. 2012. Delineation of joint molecule resolution pathways in meiosis identifies a crossover-specific resolvase. Cell 149: 334–347.

Zeng X, Saunders WS. 2000. The Saccharomyces cerevisiae centromere protein Slk19p is required for two successive divisions during meiosis. Genetics 155: 577–587.

Zhou X, Li W, Liu Y, Amon A. 2021. Cross-compartment signal propagation in the mitotic exit network. Elife 10.

